# Human lymphoid-neutrophil/monocyte restriction co-ordinately activates increased proliferation despite parallel heterogeneity in transcriptional changes

**DOI:** 10.1101/2024.02.18.580894

**Authors:** Fangwu Wang, Laura Gonzalez, Colin Hammond, Martin Hirst, Benjamin D. Simons, Connie Eaves

## Abstract

Recent studies indicate the human lympho-myeloid restriction process to be a different and more heterogeneous one than historically inferred. Here we describe the development of bulk and clonal culture systems that efficiently support early B-lymphoid differentiation and their use to identify biological and molecular changes that accompany their initial restriction from subsets of CD34+ human cord blood cells with lympho-myeloid-limited potential. Analyses of the changes observed revealed the acquisition of B-lymphoid- and neutrophil/monocyte (NM)-restricted properties are accompanied by a concomitantly accelerated and lineage-shared cell cycling activity and loss of self-renewal properties. Parallel, single-cell transcriptome analysis identified reduced expression of multiple self-renewal-associated genes and an accompanying heterogeneous activation of lineage-regulatory modules during the production of B, NM and dendritic cell precursors. These results uncover a connected regulation of lineage-shared proliferation control with persistent heterogeneity in the biological and transcriptional changes in the same cells undergoing B and NM lineage restriction.

## Introduction

Mature neutrophils (N) and monocytes (M) as well as different types of lymphoid cells (L) have highly specialized distinct functions that are essential components of a fully competent immune system. These mature blood cells are now known to be acquired through successive molecular changes that start with a loss of the long-term multi-lineage self-renewal potential of hematopoietic stem cells (HSCs) followed by a succession of changes that progressively limit the lineage and proliferative potential of their progeny^1,2^. Current findings suggest that human L and NM lineages are generated from a common, but dually restricted, CD34+ progenitor type recently identified phenotypically as LMPP (CD38-)^3^ and P-mix (CD38^med^)^4^. Downstream phenotypes with increasingly L-restricted^5–8^ or NM-restricted^9–11^ activities have also been recently reported. However, the order, coordination and pace of events that lead to the acquisition of such L- and NM-restricted phenotypes have remained poorly understood.

In contrast, homeostatic production of cells of the erythroid blood cell lineage is well established to involve a high degree of sequential coordinated changes in cell cycle, survival and apoptosis properties^12,13^. Accumulating evidence also points to important roles of external factors in eliciting differential responses of the cellular activities of different primitive progenitor subsets^4,14–17^. For B lineage differentiation, the most rapid cell expansion occurs during the assembly of the Ig heavy chain in pro-B and pre-B cells^18–20^. However, differentiated naive B cells also remain susceptible to clonal expansion in response to antigenic stimulation in secondary lymphoid organs. Previous studies also suggest that L and NM lineage restriction is largely determined by the sequential activation and interactions of several TFs, such as PU.1, IRF8, TCF3 and EBF1^21–25^. However, recent single-cell omic studies also suggest a more complex process characterized by a continuum of heterogeneously changing cell states as reflected in their gene expression programs^26–28^. In addition, dynamic changes in epigenetic modifications and chromatin organization are becoming increasingly recognized as important features of the spatio-temporal gene expression changes seen in differentiating hematopoietic cells^29–32^.

To interrogate more deeply the nature and regulation of human L+NM bi-potent cells and the generation of their derivative L- and NM-precursors, we conducted biological and molecular analyses of bi-potent (B+NM) human cells undergoing restrictive changes. The P-mix phenotype of CD34+ cord blood (CB) cells was chosen as a starting population given its high enrichment in progenitors with dually B+NM-restricted activities and ability to produce restricted progeny within 2-3 weeks in cultures containing MS5 stromal cells^4^. Further optimization of this culture system enabled us to track the initial as well as later divisions of P-mix cells with improved sensitivity and specificity. This in turn led to the discovery of several novel features of this process as described below.

## Results

### CD45RA and CLEC12A phenotypes distinguish early stages of B and NM progenitor restriction

To develop an experimental system that would allow a timed analysis of the early stages of lympho-myeloid restriction, we first measured the outputs of P-mix cells in a liquid culture system that supports the generation of both CD19+ (B) as well as CD15+ (N) and CD14+ (M) cells over a 3-week period (Fig. 1a). FACS analysis of the cells generated in these cultures revealed the rapid emergence in the first week and subsequent exponential-like expansion, of cells expressing distinguishing levels of the surface markers CD45RA (RA) and CLEC12A (C) (Fig. 1b, c). Notably, CD19+ (B-lineage) cells were subsequently detected almost exclusively in the RA+C-subset, and CD14+/CD15+ (NM-lineage) cells primarily in the C+ cells (Fig. 1b). In addition, high numbers of Lin-cells were observed in each of the RA-C-, RA+C- and C+ subsets (Fig. 1c). To enable clonal tracking of these changes, we then deployed a modified 2-step stromal co-culture procedure in which the RA-C-, RA+C- and C+ subsets of CD34+ cells present at the end of the first week (Fig. 1d, e, Supplemental Fig. S1a) were then individually assessed for their content of CD19 (B) and CD14/15 (NM) cells. This experiment showed the majority of the B+NM bi-lineage outputs, and all of the rare “blast” colonies, were obtained from RA-C-input cells (Fig. 1f). In contrast, the RA+C- and C+ input cells produced mostly pure B- and pure NM-outputs, respectively, suggesting the initiation of restriction towards these two lineages. RT-qPCR analysis of the expression of selected genes historically associated with B and NM differentiation^33,34^ in the week 2 output subsets showed a generally higher differential expression of both early and later stages of B-lineage genes (i.e., *DNTT, RAG1, CD79A, IGLL1, EBF1* and *PAX5*) in CD34+/-RA+C-cells (Fig. 1g). In contrast, the C+ phenotypes exhibited increased expression of N- and M-associated genes (i.e., *CSF1R, MPO, FCGR1A, SPI1, CEBPA*). These qPCR results point to a strong, albeit incomplete, respective association of the RA and C phenotypes with B and NM restriction.

**Figure 1.**
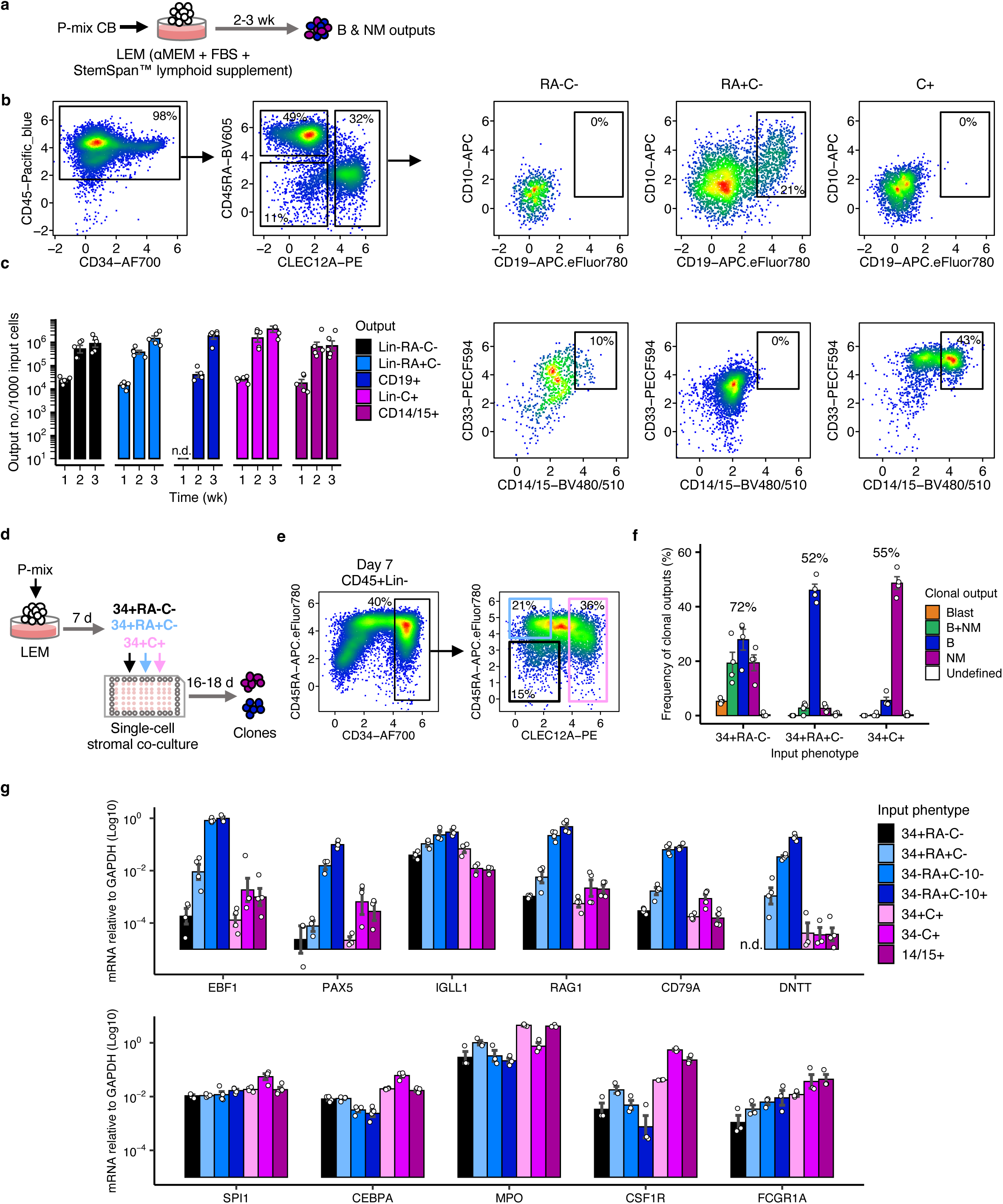
New phenotypes of cells at early stages of B and NM differentiation. **(a)** Experimental design used to generate phenotypically defined CD19+ (B) and CD14/15+ (NM) outputs *in vitro* from 1,000 P-mix CB input cells in LEM cultures (LEM being composed of αMEM, FBS and a StemSpan lymphoid expansion supplement – see Methods for additional details). **(b)** A representative flow cytometric profile of the surface marker expression of the cells present in the cultures described in (**a**) after 2 weeks. **(c)** Number of output cells per 1,000 input P-mix cells analyzed weekly. The Lin-phenotypes were gated within the CD45+14-15-10-19-subset. Each bar shows the mean ± SEM of values pooled from 5 different experiments. n.d. represents signals below the limit of detection (10 cells). **(d)** Experimental design used to examine the lineage potentials of the 3 input phenotypes shown. **(e)** FACS gating used to identify 3 phenotypes within the CD34+ cells present in day 7 cultures of CD45+14-15-10-cells generated from P-mix CB cells. **(f)** Percentages of output cell types (identified by their phenotypes) obtained in clonal cultures initiated with the 3 phenotypes shown in (**e)**. Each bar shows the mean ± SEM of values pooled from 4 different experiments with 150-350 total single cells tested per input subset. Numbers above the bars denote the overall clonal output frequencies. **(g)** RT-qPCR analysis of mRNAs of various historically defined B-lineage (upper) and NM-lineage (lower) associated genes measured in sorted phenotypes generated from 2-week cultures initiated with P-mix CB cells. Each bar shows the mean ± SEM of values pooled from 3-4 different experiments. n.d. denotes signals below the limit of detection (PCR cycle >40).

We then asked whether the phenotype changes observed in our cultures recapitulate processes associated with the generation of B and NM cells *in vivo*. To assess this, the RA-C-, RA+C- and C+ phenotypes were isolated directly from CD34+ CB cells and CD34+ CB cell-derived xenografts. To enrich for early B and NM progenitors in the CB samples, we first excluded phenotypes previously shown to be highly restricted to the erythroid, NM and B lineages (i.e., P-E, P-NM, and P-L phenotypes^4^) (Fig. 2a). Clonal outputs from RA-C-, RA+C- and C+ subsets present in the remaining CD34+38+ fraction were then assessed using the same 2-step culture method shown in Fig. 1d. The results showed highly consistent output phenotypes and at similar frequencies to those obtained from the culture-derived input cells (Fig. 2b), albeit with higher frequencies of cells with exclusively NM clonal outputs from the RA+C-as well as the RA-C-input cells.

**Figure 2.**
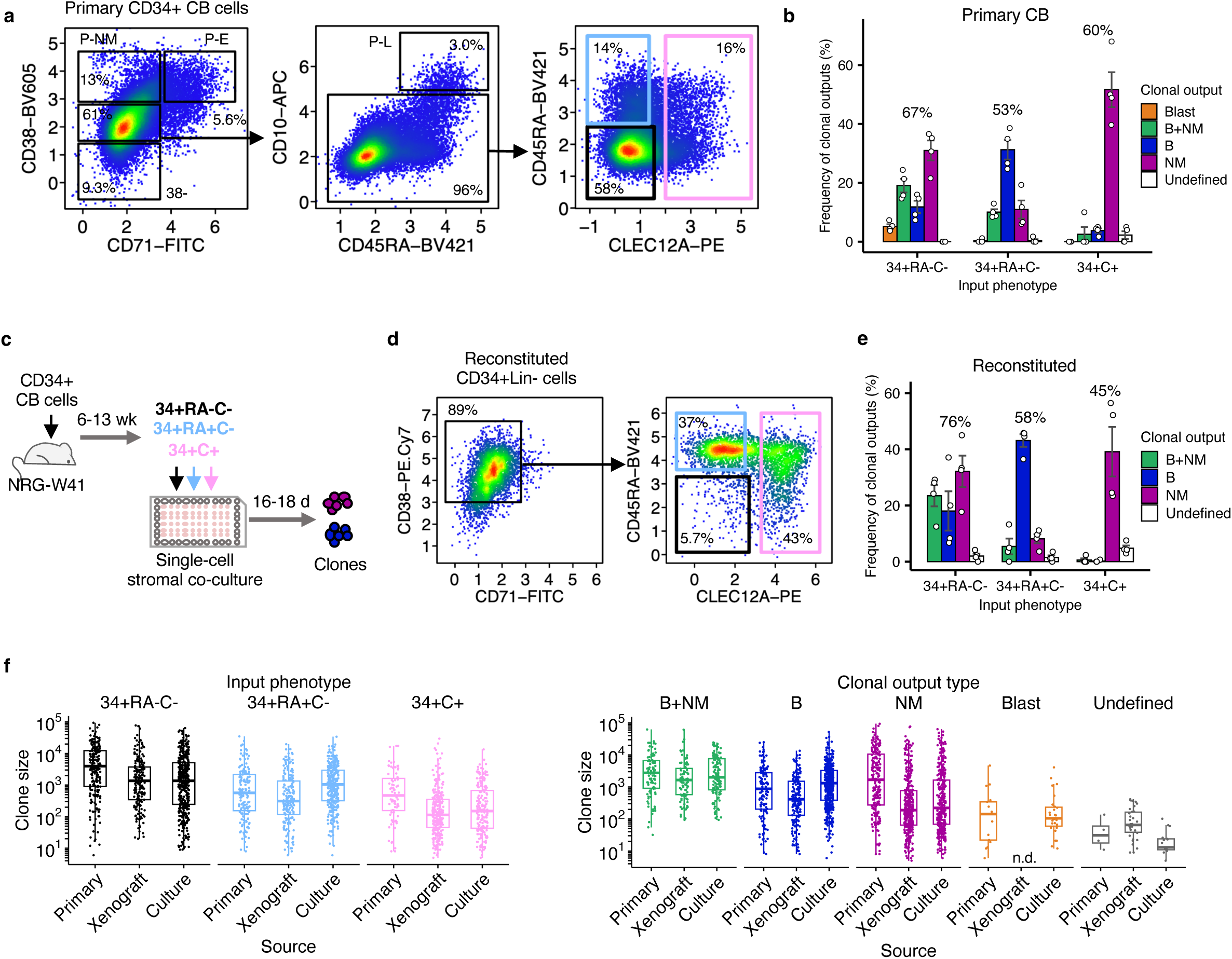
Preservation of *in vitro* determined early B and NM precursor phenotypes in CB-derived xenografts. **(a)** FACS gating used to define RA-C-, RA+C- and C+ phenotypes in unmanipulated CD34+38med71-10-CB cells. **(b)** Clonal output frequencies of the 3 input phenotypes isolated from unmanipulated CD34+ CB cells. Each bar shows the mean ± SEM of values pooled from 4 different experiments with 150-340 single cells tested per input subset. Numbers above the bars denote the overall clonal output frequencies. **(c)** Experimental design used to analyze clonal outputs of RA-C-, RA+C- and C+ phenotypes obtained from the BM of NRG-W41 mice transplanted with CD34+ CB cells (4 experiments). **(d)** FACS gating used to detect the phenotypes generated in (**c)**. **(e)** Clonal output frequencies of 3 input phenotypes isolated from the BM of engrafted NRG-W41 mice. Each bar shows the mean ± SEM of values pooled from all experiments with 260-550 total single cells tested per input subset. Numbers above the bars denote the overall clonal output frequencies. **(f)** Size of clones generated from single input cells isolated from the different sources described above. Clones (defined as ≥5 human cells) pooled from all experiments shown above were grouped by the input phenotype (left) or lineage output type (right). n.d. represents signals below the limit of detection (5 cells).

To assess the B and NM precursor outputs from CD34+ CB cells generated in engrafted sublethally irradiated NRG-W41 mice (initially injected IV 6-13 weeks previously with 2-10 x 10^3^ CD34+38-CB cells, Fig. 2c), the human CD34+RA-C-, CD34+RA+C- and CD34+C+ cells present were then isolated and clonally analyzed *in vitro* for their output capabilities (again as shown in Fig. 1d). The results showed similar clonogenic frequencies and lineage outputs as seen for their counterparts from the original culture system and primary CB analyses, although in this latter case, detectable levels of blast colonies were not observed (Fig. 2d, e). The final clonal output numbers were also highly variable among clones with shared input phenotypes or lineage output types regardless of the initial source of input (Fig. 2f). Additional analyses of directly isolated human adult BM CD34+ cells showed that they also contain the same RA and C phenotypes (Supplemental Fig. S1b). Notably, the variabilities in the relative frequencies of the RA and C phenotypes from the same initial sources were less pronounced than between the different sources analyzed (Supplemental Fig. S1c). Taken together, these results suggest the early steps of B and NM restriction may be associated with consistent initial phenotypic changes.

In addition to examining B- and NM-lineage outputs, we found that the original P-mix input cells also contained cells capable of generating CD33+CD1A+ dendritic cells (DC), and CD45RA+CD123+ plasmacytoid DC lineages (Fig. 2i, S2b), whereas the CD34-RA-C-output cells were found to also give rise to glycophorin A+ (erythroid) cells in a secondary culture supplied with EPO and a different growth factor (GF) combination.

### B and NM restriction processes are accompanied by concurrently enhanced proliferative activity

Successive increases in the proliferative activity of hematopoietic cell progenitor populations as they undergo separated erythro-myeloid differentiation processes is well documented^35–37^. However, if and when a similar change might begin in cells undergoing lympho-myeloid differentiation is unknown. To address this question, we first generated CD34+RA-C-cells from P-mix CB cells in 1-week suspension cultures and then labelled them with Carboxyfluorescein succinimidyl ester (CFSE) prior to using them to initiate secondary cultures using the same culture conditions (Fig. 3a). Subsequent flow cytometric tracking of the CSFE-labelled peaks obtained over the next 8 days showed the size of the subpopulation maintaining a CD34+RA-C-phenotype increased only slightly (∼3-fold) by day 8 (Fig. 3b), whereas the differentiated RA+C- and C+ progeny populations expanded exponentially (∼500-fold) during the same period.

**Figure 3.**
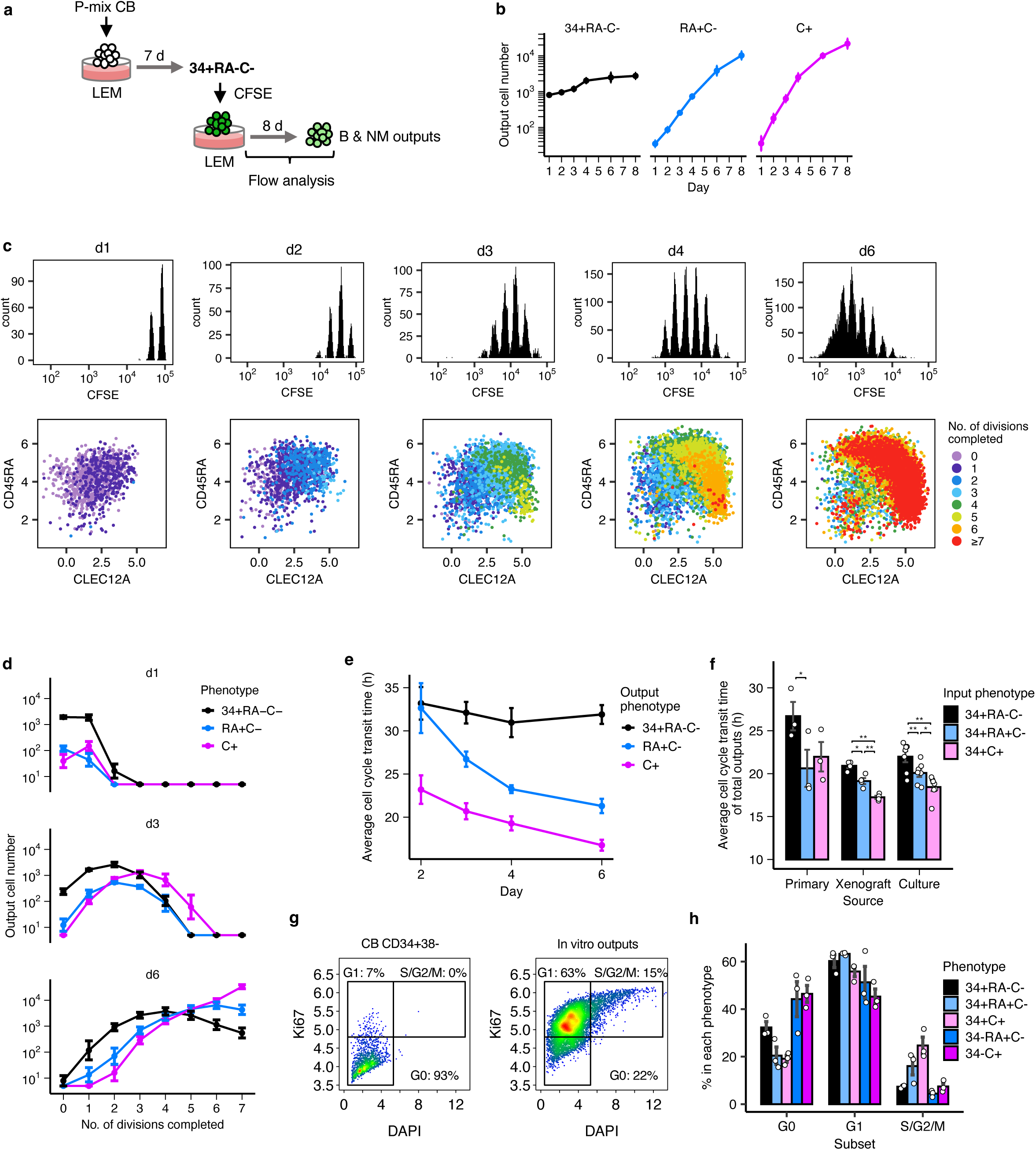
Cells differentiating toward either B or NM lineages undergo rapid and simultaneous acceleration of their proliferative activity. **(a)** Design of the cell division tracking strategy used to analyze the progeny of individual CD34+RA-C-cells. **(b)** Number of output cells per 1,000 input CD34+RA-C-cells tracked over an 8-day period in the design shown in (**a)** (data points show the mean ± SEM values pooled from 5 experiments). **(c)** CFSE and surface marker expression profiles of cells obtained at different times after initiating cultures of CFSE-labelled cells. The number of completed divisions was determined by the fold-dilution of CFSE fluorescent intensity. **(d)** Numbers of the 3 output phenotypes detected in successive CFSE fluorescent peaks indicative of completion of different numbers of cell divisions (generated per 1,000 CD34+RA-C-input cells; data points showing the mean ± SEM values pooled from 5 experiments). **(e)** Average cell cycle transit times (in hours) preceding the appearance of different output phenotypes at different time points (data points showing the mean ± SEM values pooled from 5 experiments). **(f)** Average cell cycle transit times (mean ± SEM) of the immediate progeny of different sources of CFSE-labelled CD34+RA-C-, CD34+RA+C- and CD34+C+ phenotypes assessed after a 4-day culture in LEM. Input cells tested were isolated directly from unmanipulated CD34+ CB cells (n=3 experiments), from xenografts (n=4 experiments) and cultures (n=7 experiments). *p<0.05, **p<0.01 via pairwise t-tests post Holm adjustment. **(g)** Assignment of G0, G1, S/G2/M phases based on the intensity of Ki67 (log10-transformed pixel) and DAPI (10^5^ pixel) measured by immunofluorescence. Data shown are primary CB CD34+38-cells (left) and pooled output populations from one experiment. **(h)** Proportions of cells at each cell cycle phase (mean ± SEM) within each phenotypic population (n=3 experiments).

To compare the lengths of the cell cycle associated with these two distinct patterns of cell population expansion, the division histories of the output cells were determined quantitatively from the sequential dilution of their retained CFSE signal (Fig. 3c). The results showed a trend towards a positive correlation between their RA or C surface marker expression and the accumulated number of cell divisions (Fig. 3c, d). In contrast, maintenance of the CD34+RA-C-phenotype was clearly associated with the completion of fewer cell cycles within the same timeframe (Fig. 3c, d). Based on this observation, average cell cycle lengths were calculated according to the distribution of cell division numbers at different time points (see detailed calculation method in Supplemental Fig. S3). This revealed a shortened cell cycle in both RA+C- and C+ output phenotypes already by day 6 (21 h and 17 h) compared to the input CD34+RA-C-cells (32 h) (Fig. 3e), indicating a marked increase in cell division times accompanying the production of the RA+C- and C+ cells from their RA-C-precursors.

To determine whether these differences in cell cycle behaviour would accompany the generation *in vivo* of the same cell phenotypes, different CD34+ subsets isolated directly from CB, and from cells regenerated in CD34+ CB-transplanted NRG-W41 mice were similarly labelled with CFSE and then examined *in vitro* (Fig. 3f). The average division intervals in cells from these sources when assessed after 4 days *in vitro* showed similarly more prolonged cell cycle transit times of the outputs of RA-C-cells than for the outputs of the RA+C- and C+ cells. In addition, the pace of cell divisions exhibited by the same phenotypes isolated directly from CB were comparable to their counterparts generated in culture and in the xenografts, suggesting a consistent pattern of activation of proliferation activities accompanying the onset of lymphoid and myeloid differentiation in these different settings.

These results prompted us to further compare the distribution of cells at different cell cycle phases. Examination of the intracellular levels of Ki67 and DNA content in individual cells using primary CD34+CD38-cells as a “G0” control (Fig. 3g, h) showed only 68% of CD34+RA-C-cells to be in cycle, with an initial increase in the fraction of cycling cells in the CD34+RA+C- and CD34+C+ subsets (80% and 81%). In contrast, the proportion of non-cycling cells was again increased in the CD34-subsets, suggesting a partial loss of proliferative activity in their more differentiated progeny.

The CFSE data also showed heterogeneity in the number of divisions involved in the generation of the CD34+RA+C- and CD34+C+ cells. To examine this at a clonal level, the number and phenotype of outputs of single CD34+RA-C-*in situ*-labelled input cells were tracked in 16-day clonal cultures (Fig. 4a, b) and output phenotypes determined by the subsequent appearance of different positive cell phenotypes in cultures preloaded with fluorescent antibodies against CD34, RA and C epitopes. This revealed 1 in 5 input cells produced bi-lineage B+NM cells (Fig. 4c). Moreover, retrospective tracking of the phenotypic changes in these bi-lineage clones showed that half of them displayed exclusive outputs of unchanged CD34+RA-C-phenotypes for the first 4 days (indicative of “self-renewal” divisions). However, by day 8, most of these clones had produced cells with one or both differentiated phenotypes (i.e., CD34+RA+C- and/or CD34+C+ outputs, Fig. 4d). Sub-classification of the clones according to the timing of a first appearance of differentiated phenotypes revealed distinct clonal expansion kinetics among them (see pooled average and individual clone data in Fig. 4e and Supplemental Fig. S4, respectively). For example, clones in which differentiated phenotypes appeared within 6 days showed initially higher rates of expansion that then decreased in comparison to clones with more delayed evidence of differentiation (>6 days). Empirically, the pooled average clone data fit well with a logistic growth dynamic (Fig. 4f), while measurements of the median lengths of the delay in accelerated proliferation (λ) across clones were found to be roughly proportional to the delay in differentiation to a restricted phenotype (Fig. 4g). This association suggests that the increase in proliferation rate is strongly linked to the transition from a phase of slow CD34+RA-C-self-renewing divisions to the initiation of a more rapid lineage restrictive mode of cell proliferation. In addition, this analysis showed that the maximum rate of clone expansion (μ) and its final output (A) were both higher in clones with evidence of a later onset of restriction (Fig, 4g).

**Figure 4.**
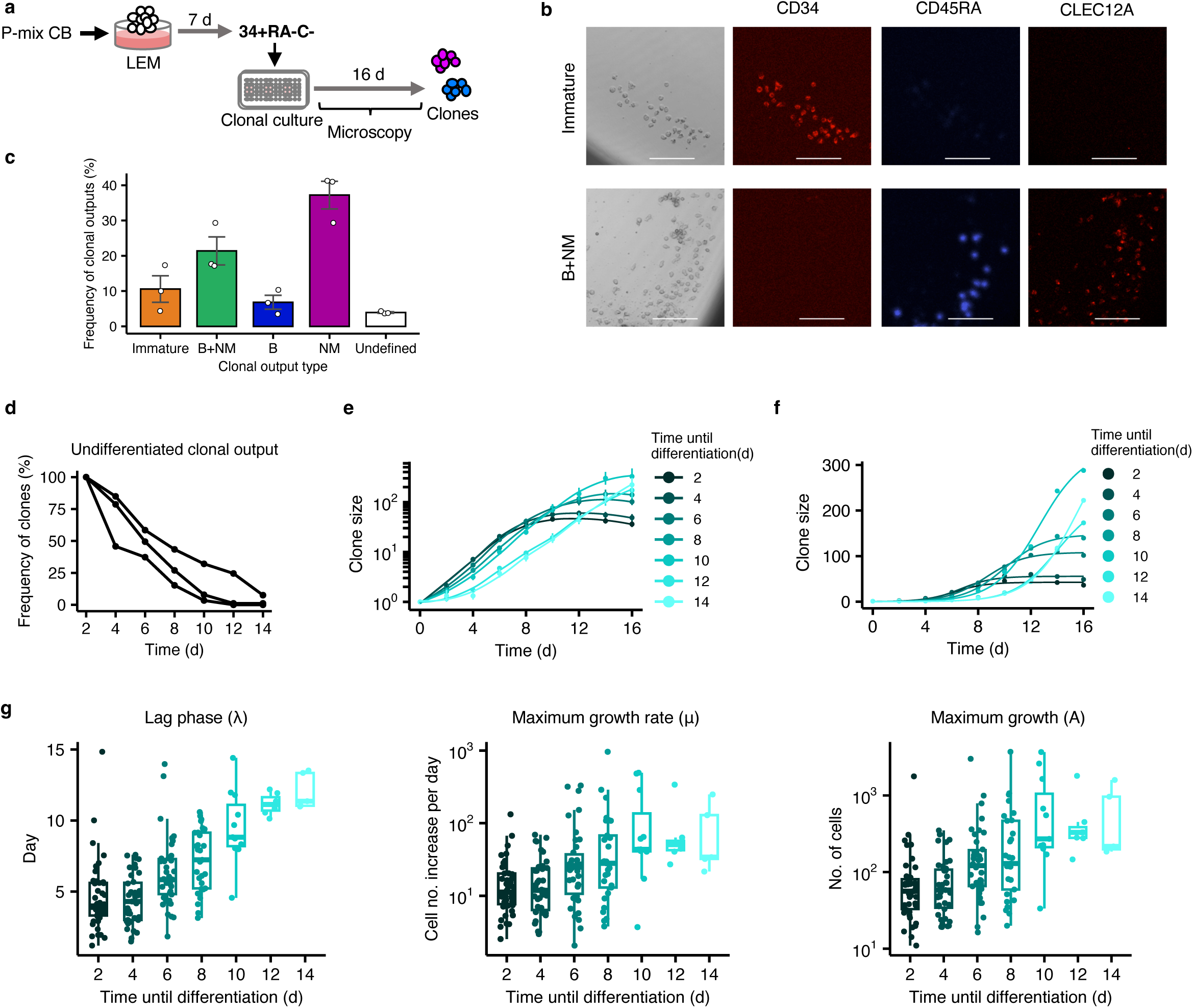
Clonally tracked B+NM progenitor outputs reveal initial slow divisions are associated with larger cell outputs. **(a)** Experimental design used to track clonal changes in the surface expression of CD34, CD45RA and CLEC12A. **(b)** Bright-field and fluorescent images of representative clones examined on day 12. Scale bar = 100 μm. **(c)** Percentages of clones with different 16-day output phenotypes obtained from single CD34+RA-C-input cells. Each bar shows the mean ± SEM of values pooled from 3 experiments with 960 total single cells examined. **(d)** Percentages of clones that initially produced CD34+RA-C-outputs exclusively at different time points, but later produced RA+C- and/or RA-C+ cells. Each line represents the results from each such experiment performed. **(e)** Cumulative growth kinetics (Loess-smoothened) of clones initiated from CD34+RA-C-cells (n=201) according to the time of the first appearance of RA+C- or C+ cells in them. **(f)** Growth-curves of the geometric means of the clone numbers analyzed in **e** and generated using a logistic model. **(g)** Growth parameters of individual clones analyzed in **d** and **e**. Each data point represents a clone (center line=median; box limits=first and third quartiles; whiskers=1.5x interquartile range).

To further assess how precursor proliferation rates may be differentially linked to lineage restriction probabilities, we developed a strategy to prospectively isolate input cells with different cycling properties. To this end, CD34+RA-C-cells were first isolated from the outputs of a 1-week culture initiated with P-mix CB cells (Fig. 5a). They were then labeled with CFSE, cultured for 4-6 days in the liquid culture condition and their outputs then isolated by FACS according to their preceding slow- or fast-cycling history. The subsequent outputs of these 2 cohorts were then separately assessed further in 6-day cultures. The results show that the more slowly dividing input cells generated higher CD34+RA-C-outputs than the more rapidly dividing input cells and the latter also produced higher outputs of RA+C- and C+ progeny (Fig. 5b). Assessment of the speed of acquisition of B and NM differentiation properties by similarly isolated slow and fast-cycling CD34+RA-C-cells was then also assessed in the clonal culture system shown in Fig. 1d (Fig. 5c). This further revealed that the more slowly dividing input cells generated more dual lineage (B+NM) clones and fewer restricted (NM) clones compared to the initially more rapidly dividing input cells (Fig. 5d). In addition, clones composed of CD34+ blasts were detected only in clones generated from the slow-cycling input cells. Taken together, these results provide further support for the concept that CD34+RA-C-cells with dual B+NM potential lose their ability to maintain this state as they acquire shorter cell cycle transit times.

**Figure 5.**
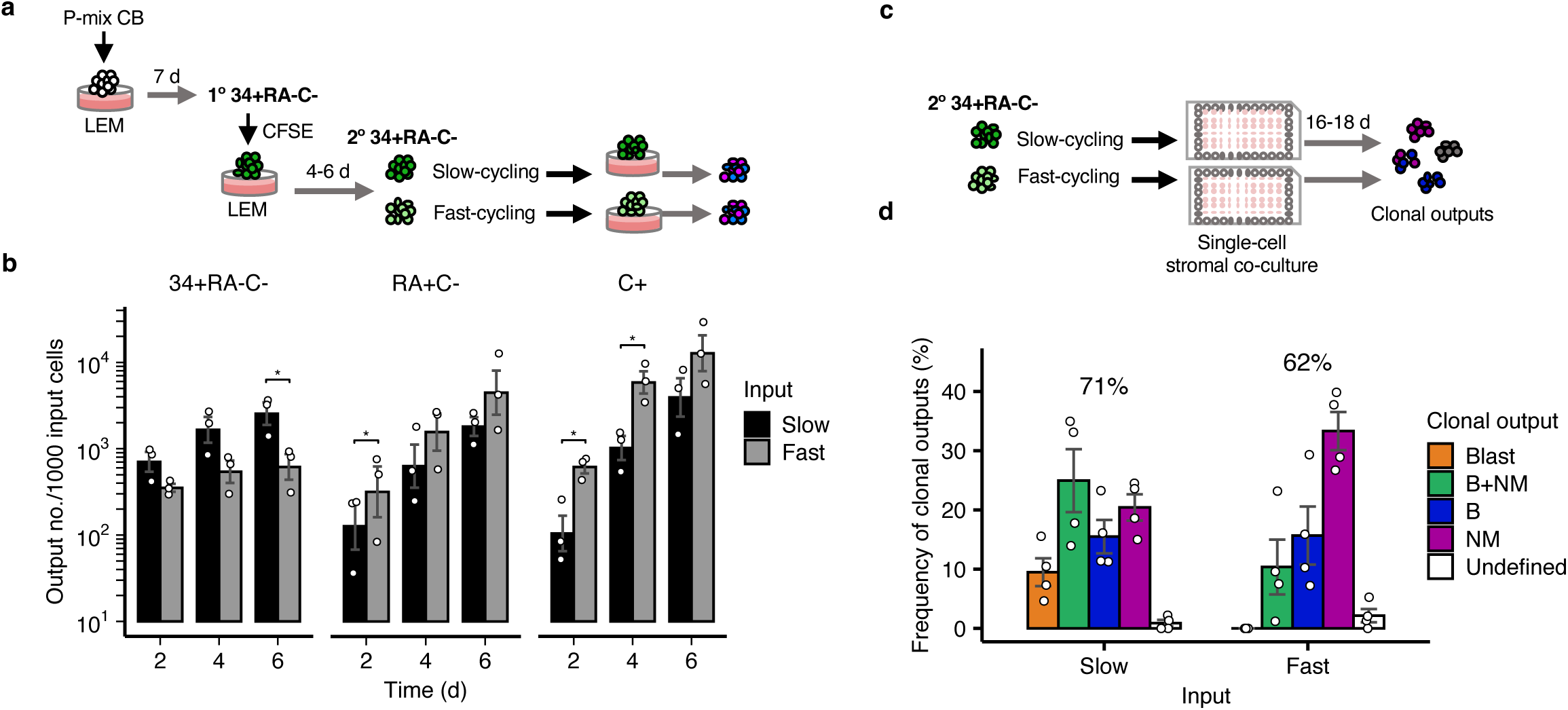
Evidence of a delayed B/NM differentiation process in CD34+RA-C-cells exhibiting a slower cycling behaviour. **(a)** Experimental design used to compare the changing output phenotypes generated from slow- and fast-cycling CD34+RA-C-cells. **(b)** Output numbers (mean ± SEM) of different progeny phenotypes generated from slow- and fast-cycling input CD34+RA-C-cells determined at different time points (n=3 experiments). *p<0.05 via pairwise t-tests post Holm adjustment. **(c)** Experimental design used to examine and compare the clonal outputs of individual slow- and fast-cycling CD34+RA-C-cells. **(d)** Percentages of clonal output types produced from CD34+RA-C-input cells. Each bar shows the mean ± SEM of values pooled from 4 different experiments in which a total of 540 and 520 single cells were assessed from the slow- and fast-cycling progenitors, respectively. Numbers above the bars denote the overall clonal output frequencies.

### Self-renewal-associated genes are highly expressed in less proliferative CD34+RA-C-cells

To analyze the changes in gene expression that accompany the differentiation of CD34+RA-C-cells, we applied CITE-seq analysis to cells harvested in the 2-step culture protocol described in Fig. 6a. Simultaneous detection of antibody-derived nucleotide signals (ADTs) enabled the transcriptome data to be assigned to particular phenotypes using nucleotide-linked antibodies to CD34, CD45RA, CLEC12A, CD10, CD7, CD14 and CD15. Initial analysis of cell cycle changes showed nearly 100% of the input P-mix CB (“d0”) cells could be readily classified as G0/G1 cells (Fig. 6b, lower panel). Evidence of high cycling activity became apparent on day 10 and then appeared decreased on day 13. Analysis of other transcriptional changes after cell cycle regression yielded a UMAP with a clear segregation of 2 clusters in the first dimension (Fig. 6c). Genes highly expressed in the smaller Cluster (B) were identified as erythrocyte and megakaryocyte-associated (e.g., GATA1, GATA2 and HBD), and genes expressed at higher levels in Cluster A were associated with lymphocyte and leukocyte differentiation (Supplemental Fig. S5a, b, Supplemental Table 5). More detailed examination of the transcripts associated with the Cluster A cells in the 2nd and 3rd dimensions of the UMAP revealed a continuum of output cells progressing from the right to the left of the time course analysis (Fig. 6d). Mapping of the CITE-seq signals to established surface marker phenotypes revealed a gradual shift from early to differentiated phenotypes (Fig. 6e). However, these did not show distinct separate groupings. To assess directly the relationship between cell cycle and transcriptional states, cells were mapped to their assigned cell cycle phases (Fig. 6f). This revealed a relatively high proportion of CD34+RA-C-cells, as well as their more restricted CD34-RA+C- and 14/15+ cells, were in G0/G1. In contrast, the majority of the intermediate precursors appeared to be actively cycling (Fig. 6g).

**Figure 6.**
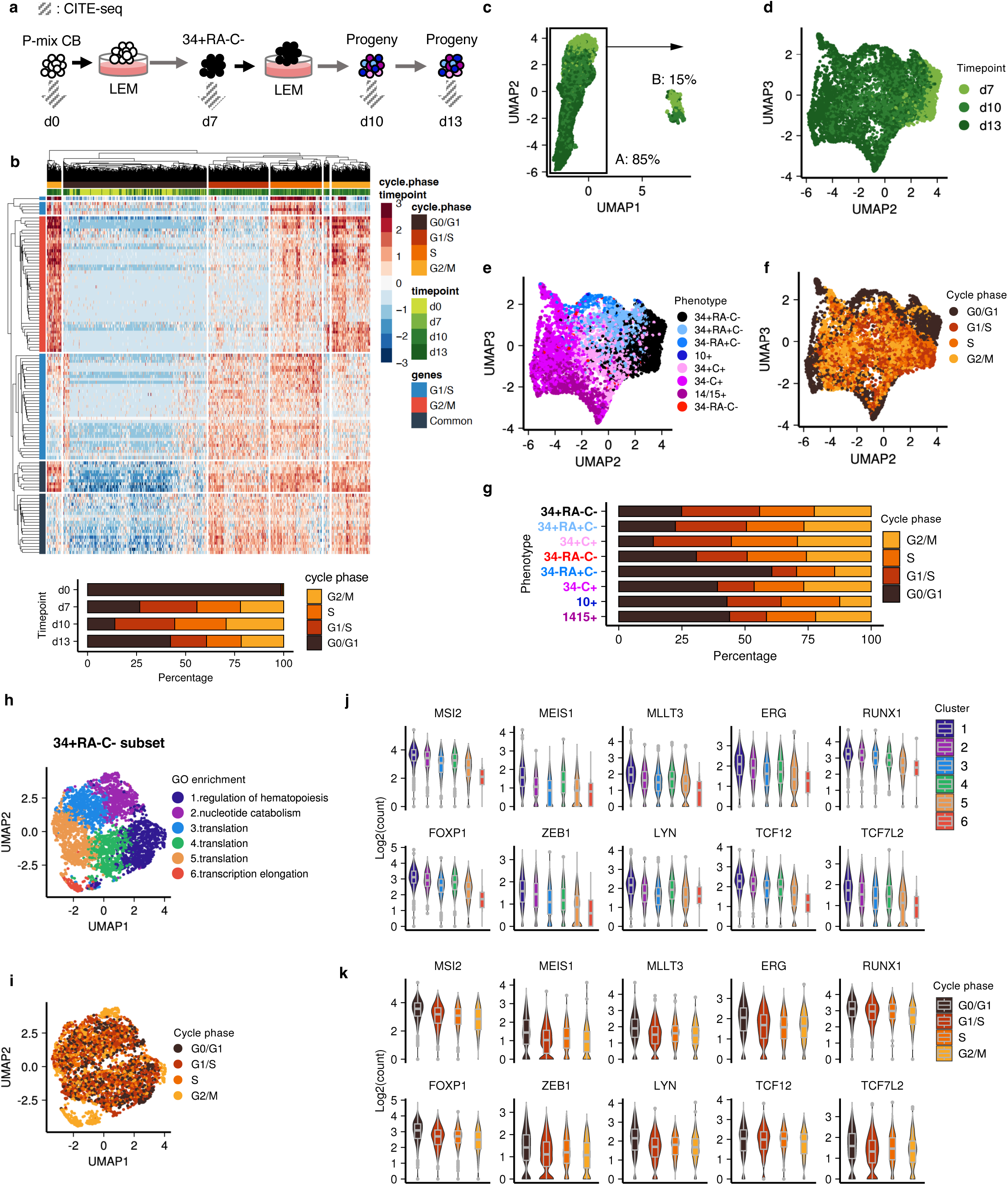
CD34+RA-C-cells in a lower proliferative state display higher self-renewal transcriptional programs when stimulated to differentiate. **(a)** Experimental design used to track the sequential transcript outputs of *in vitro*-stimulated P-mix CB cells by single-cell CITE-seq analysis. For details, see Methods. **(b)** Assignment of cell cycle phases based on the expression of cell cycle-associated genes. In the upper panel, the columns represent 16,269 individual cells coloured by time point and cell cycle states. The rows represent genes specifically expressed in certain cell cycle phases. Cell colours on the heatmap denote the z-score of log-transformed gene expression values. The percentages of cells in different cell cycle phases at each time point assessed are shown in the lower panel. **(c)** UMAP presentation (post cell cycle regression analysis) of transcriptome data combined from 12,181 single cells from the day 7, 10 and 13 time points. Cells were divided into “A” and “B” clusters based on the first UMAP dimension. (**d**-**f**) UMAP projection of 10,385 cells in the “A” cluster. Individual cells were coloured by time (**d**), cell cycle phase (**e**), or surface phenotype inferred from antibody-derived oligonucleotide tag signals (**f**). (**g**) Comparison of the distribution of different cell cycle phases in Cluster A cells separated by different phenotypic subsets. (**h**-**i**) UMAP display of the clustering (**h**) and cell cycle status (**i**) of the phenotypically defined CD34+RA-C-subset in Cluster A cells. Each cluster in (**h)** is annotated by the highest-ranked GO term (q-value cutoff <0.05). (**j**-**k**) Log-transformed transcript counts of genes ranked highly in subcluster 1. Each point denotes a cell colored by its designated subcluster (**j**) or cell cycle phase (**k**) (center line=median; box limits=first and third quartiles; whiskers=1.5x interquartile range; points=outliers).

We then asked whether specific transcriptional patterns could be associated with initial changes in cell cycle states. Accordingly, the subset of immature CD34+RA-C-cells pooled from all time points were first grouped into different clusters and the most enriched features of each cluster then identified based on their differential gene expression (Fig. 6h, i). Interestingly, genes highly expressed in Cluster 1 were enriched for TFs associated with primitive hematopoietic cell functions, such as MEIS1^38^, MLLT3^39^, HMGA2^40^ and MSI2^41^, and those in Cluster 2 were enriched in transcripts of genes implicated in lymphoid cell regulation, such as FOXP1, ZEB1 and the E proteins^42–44^ (Fig. 6j, Supplemental Table 6). In addition, these genes were also found to be expressed at higher levels in the G0/G1 cells compared to cells in the other cell cycle phases (Fig. 6k). Conversely, Clusters 3-6 were highly enriched for transcription and translation activities. Interestingly, genes encoding proteins previously shown to mediate cell type-specific or signal transducing functions were rarely detected in CD34+RA-C-cells irrespective of their cell cycle states (Supplemental Fig. S6a).

### Sequential activation of differentiation regulator transcripts accompanies B and NM lineage restriction

Application of an unsupervised lineage analysis to the transcriptome data identified 4 trajectories that all started from a cluster overlapping with those of the CD34+RA-C-population (Fig. 7a). Based on their historically recognized lineage-associations, we called these paths-B, -DC, -M and -N, respectively (Fig. 7b, Supplemental Fig. S6b). Dynamically expressed genes were then classified as early, intermediate or late according to the pseudo-timing of their most active expression in a particular lineage (Fig. 7c). Intersection of these data showed the majority of lineage-specific genes were activated late (Fig. 7d, Supplemental Fig. S6c), whereas the early and intermediate genes showed extensive overlap between lineages. However, a group of genes including TF genes associated with primitive CD34+RA-C-cells (Fig. 6j) also showed early or intermediate expression patterns only in the M trajectory (Fig. 7e). Similarities between gene expression levels across all 4 lineages were further compared using a UMAP strategy (Fig. 7f). Gene ontology analysis showed the “late genes” overlapped with clusters enriched for the functions of the corresponding cell type, whereas the intermediate genes ubiquitously expressed in all lineages were mapped to clusters enriched for translation and nucleotide metabolism functions, consistent with the increased proliferation activities of these cells (Fig. 7g, Supplemental Table 7).

**Figure 7.**
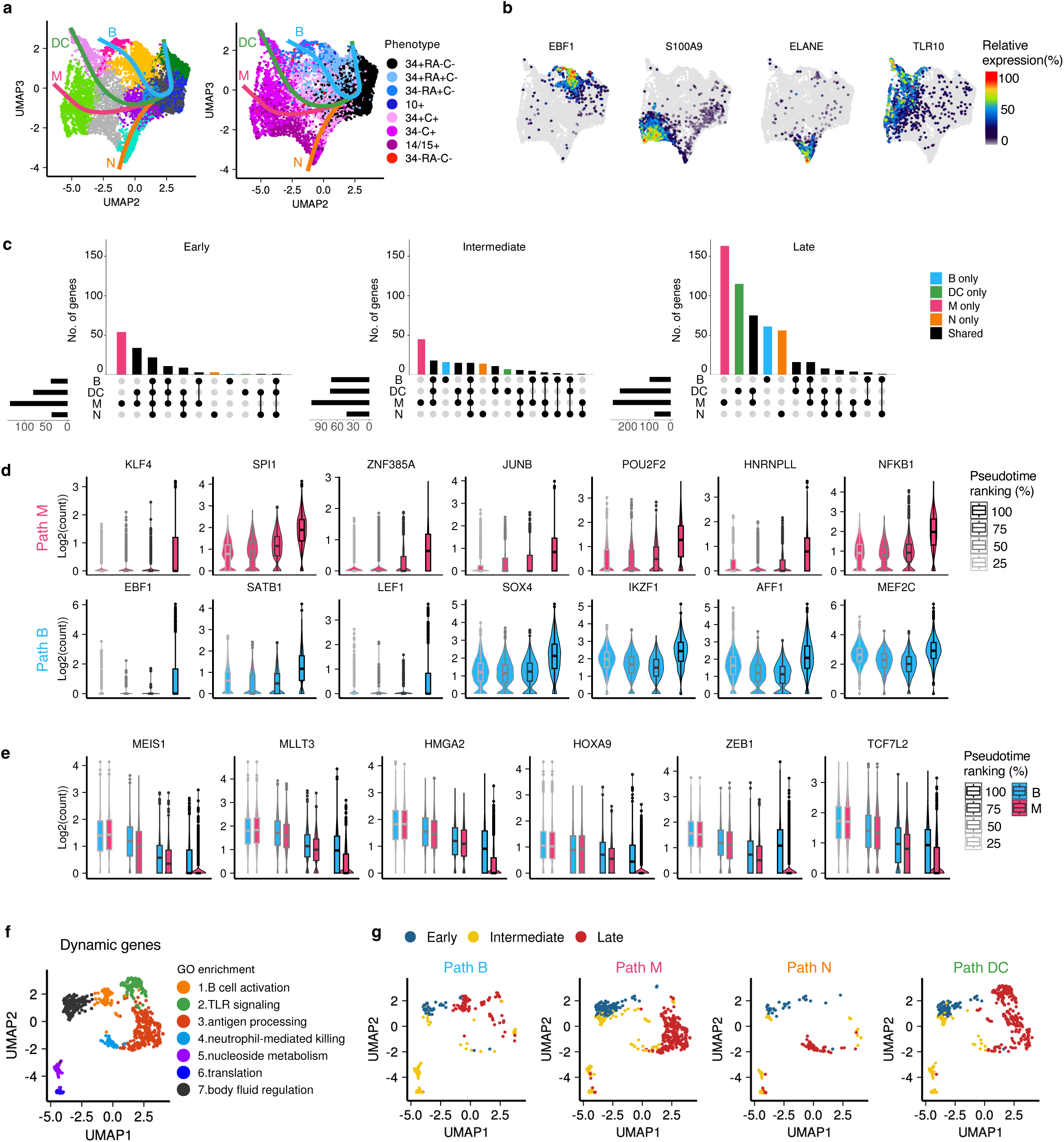
Lineage trajectories inferred from timed changes in expression of self-renewal and lineage-promoting genes. **(a)** Lineage trajectory predictions of pre-determined cell clusters embedded in the UMAP manifold. Each colored line represents a single trajectory path. Each cell is colored by its cluster designation (left) or assigned phenotypic subset (right). **(b)** Gene expression in Cluster A cells normalized to the maximum log-transformed counts of each gene representative of B, M, N and DC lineage-specific features, respectively. **(c)** Gene expression intersects of the different lineage paths that displayed early (pseudotime 0-20%), intermediate (pseudotime 20-80%) or late (pseudotime 80-100%) expression patterns. (**d**-**e**) Changes in the expression of “late” genes (**d**) and “early” genes (**e**) across the M and B trajectories. Cells are grouped by the quadrants of their pseudotime ranking (center line=median; box limits=first and third quartiles; whiskers=1.5x interquartile range; points=outliers). **(f)** UMAP presentation and nearest neighbour clustering of genes showing the most dynamic changes in expression. Each cluster is annotated by the highest-ranked GO term (q-value cutoff <0.05). **(g)** UMAP presentation of the most variably expressed genes in each trajectory. Genes are coloured by the early, intermediate or late expression patterns as defined in (**c**).

To further assess interactions between different TF genes, we adopted an agnostic approach to identify those predicted to affect gene expression in the current dataset using “SCENIC”^45^. This analysis revealed a number of self-renewal associated TF genes (e.g., HOXA9, HLF and HMGA2) in the primitive cells, MYC and E2F factors in the intermediate precursors, and lineage-associated TF motifs in the later stages of restriction (Fig. 8a). In addition, some TFs appeared to be relevant across all cell types (e.g., see overlapping TF motifs across the B, M and DC lineages in Fig. 8b). Overlapping activities of EBF1 and IRF family genes were also evident in a subset of primary P-mix cells (Supplemental Fig. S7). Taken together, these results indicate loosely coordinated changes in various sets of TF activities likely accompany changing states of lineage-restricted potentials.

**Figure 8.**
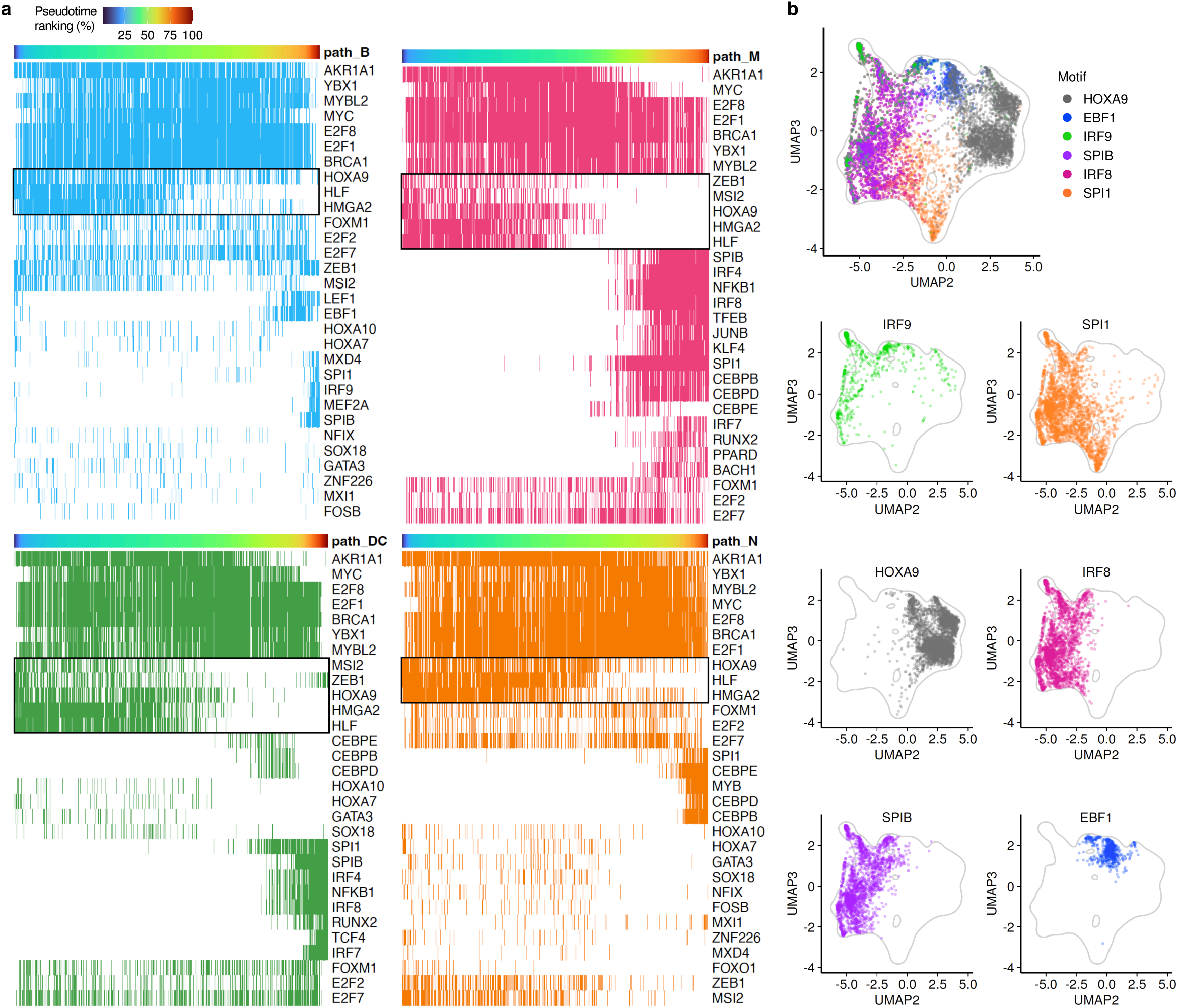
Dynamic and interactive changes in TF motifs within and across lineage restriction trajectories. **(a)** Top 30 TF motifs based on the number of cells identified in each lineage trajectory. Columns represent individual cells ordered by pseudotime. Rows represent TFs whose motifs were detected in >20 target genes. Cell colours on the heatmap denote presence (coloured by lineage) or absence (blank) of the motif activity. Black boxes denote examples of TFs associated self-renewal of early hematopoietic precursors. **(b)** Distribution of cells showing active status of each TF motif in the UMAP space of Cluster A cells. Each TF is represented by a distinct colour. Grey lines represent a density level of 0.006 of all cells in Cluster A (bin number = 100).

## Discussion

Together, these results provide new insights into the nature and related time course of changes in the lineage outputs, proliferation activities and transcriptional states of normal human hematopoietic cells with limited but still dual B+NM potential as they become further restricted. Building on previous evidence that the surface expression of CD45RA and CLEC12A strongly correlates with B and NM output potential^10,46^, the current findings reveal new evidence of an association of the differential expression of these two phenotypes with the onset of their biological restriction. Unexpectedly, our results show that in both *in vivo* and *in vitro* conditions, the restriction of precursors to the B and NM lineages is accompanied by a simultaneously timed and conserved activation of a reduced cell cycle transit time. Finally, our single cell transcriptional data provide previously undocumented heterogeneity of cell state changes that accompany early B/NM restriction events.

Interestingly, a shared activation of emerging B- and NM-restricted phenotypes is consistent with derivative lineage generation processes in other hierarchically organized tissue types, such as intestine and skin, where rapid expansion of mature functional cells occurs mainly at the level of restricted progenitors in response to extrinsic signals (GFs, cytokines, injury signals)^47,48^. Here, a lineage-shared activation of cell proliferation during B and NM restriction was shown to be transcriptionally associated with higher metabolic and translation activities but, interestingly, not strongly with various lineage-specific signaling components (e.g., IL7R, CSF1R, TLR9/10) that were rarely detected in these early precursors. However, transcripts of several genes known to be involved in the self-renewal functions of more primitive hematopoietic cells (e.g., *MLLT3*^39^*, MEIS1* and *HOXA9*^38^), were all observed in the early RA-C-precursors at higher levels than in their RA+C- and C+ progeny. This latter finding is consistent with the concept that expression of these genes may favor prolonged cell cycle transit times and perhaps greater stability of an undifferentiated state. Taken together these findings underscore the increasing importance of identifying early precursor phenotypes and tracking their clonal progeny in environments that optimize the production and characterization of their changing biological properties.

## Methods

### Primary human samples

Anonymized anticoagulated samples of umbilical CB were collected from consented parents and normal adult BM cells were collected from consented allogeneic BM transplant donors according to protocols approved by the Research Ethics Board of UBC. Fresh samples were first processed using Lymphoprep™ (STEMCELL Technologies) to separate low-density fractions from granulocytes and erythrocytes, with or without immune-based co-removal of CD11b+ myeloid cells, CD19+ B-cells and CD3+ T-cells. Positive selection of CD34+ cells was then performed on the low-density cells using the EasySep™ kit from STEMCELL Technologies) to obtain final CD34+ purities, usually of >90%. Pooled CB samples were generated by combining cells from >30 donors prior to the CD34+ selection step. Aliquoted CD34+ cell-enriched samples were cryopreserved in FBS (STEMCELL Technologies) containing 10% DMSO (Origen Biomedical). As needed, cells were rapidly thawed at 37 °C in a water bath and added dropwise into IMDM (STEMCELL Technologies) containing 10% FBS and 10 μg/mL DNase (MilliporeSigma). All *in vitro* and *in vivo* experiments were performed using individual or pooled donors and were performed according to the protocols approved by the Research Ethics Board of UBC and UBC Animal Approval and Biosafety guidelines.

### Phenotype analysis and isolation of hematopoietic cells by FACS

For bulk or clonal FACS analysis of primary cells, cells obtained from cultures, or xenografts, cells in a single-cell suspension were stained with fluorochrome-conjugated antibodies specific to surface antigens (listed in Supplemental Table 1) in Hanks’ Balanced Salt Solution (HBSS, STEMCELL Technologies) supplemented with 2% FBS, 1.5 µg/mL anti-human CD32 antibody (STEMCELL Technologies) and 10 µg/mL anti-mouse CD16/CD32 antibody (as needed) (STEMCELL Technologies) for 1 hour on ice. Cells were then washed in HBSS + 2% FBS, re-suspended in the same solution containing 1 μg/mL propidium iodide (PI, Millipore Sigma) and passed through a 40 μm filter (as needed) before being analyzed on a BD LSR Fortessa Cell Analyzer or isolated using a BD FACSAria Fusion or BD FACSAria III Cell Sorter. Correction of signal spillover for each fluorescence channel was performed using UltraComp eBeads (Invitrogen) stained with individual fluorochrome-conjugated antibodies. Hierarchical gating of cell populations were defined using forward and side light scatter (FSC and SSC) and the fluorescent signals obtained from PI and antibodies attached to surface antigens present on the surface of viable (PI-) test cells. Sorting of bulk cell subsets and single cells were performed using the “4-Way Purity” and “Single Cell” precision mode, respectively. Collected FACS data were then analyzed and plotted using FlowJo software or the R package “flowCore”. The scales of plotted surface marker parameters were transformed using the asinh (x/150) function in order to optimize visualization of the lowest and highest ranges of data on the same plot.

### LEM feeder-free cultures

The LEM condition used to generate B and NM precursors was composed of 7.5% FBS, 10% StemSpan™ Lymphoid Progenitor Expansion Supplement (STEMCELL Technologies) and 82.5% αMEM containing 2 mM glutamine and 10^−4^ M β-mercaptoethanol. Bulk cultures were initiated with P-mix or other sorted CB subsets at a density of 700-2,000 cells/100 µL. Half-medium changes were performed at the end of each week using the same media. Output cells were harvested by rinsing the wells 2 times with HBSS + 2% FBS, and then analyzed or sorted using a flow cytometer. The maximum limit of detection for each human lineage type was set at 0.005%; i.e. at least 10 cells in a minimum of 2 x 104 collected events within each category being examined. The definitions of the different lineage types assessed are listed in Supplemental Table 2.

### Mouse stromal cell lines

The MS5 stromal cell line, originally isolated and cloned from irradiated mouse bone marrow adherent cultures^49^ and obtained as gift from L. Coulombel (INSERM U362) were maintained in DMEM with 4,500 mg/L D-Glucose (STEMCELL Technologies) and 10% FBS, and passaged when full confluency was reached, typically every 3-4 days.

The M210B4 stromal cell line was isolated from normal mouse bone marrow adherent cells^50^ and maintained in RPMI 1640 medium (STEMCELL Technologies) with 10% FBS. The SL stromal cell line was isolated and cloned from *Sl/Sl* mouse embryo adherent cell cultures^51^ and maintained in DMEM with 10% FBS. Derivative M210B4 and SL cell lines transduced with retroviral vectors to produce human GFs^52^ were maintained as for the parental cell lines. All M210B4 and SL cell lines were irradiated with 80 Gy of X-rays before they were plated for co-culture experiments with human hematopoietic cells. MS5 cells were not irradiated.

### Clonal B+NM multi-lineage assays using stromal co-cultures

For clonal analysis of the B and NM outputs from single human hematopoietic cells, 9,000 MS5 cells and 330 each of the engineered M210B4 (G-CSF+IL-3), SL (SCF+IL-3) and SL (FLT3L) cell lines were plated into the 60 inner wells of a 96-well plate, and then sterile water added to the peripheral wells. The next day, individual human hematopoietic cells was sorted into each of the inner wells already preloaded with feeders and 7.5% FBS plus 92.5% αMEM containing 2 mM glutamine, 10^−4^ M β-mercaptoethanol, 50 ng/mL SCF, 10 ng/ml FLT3L and 10 ng/ml IL-7. Cultures were incubated for 3 weeks in a humidified 37 °C atmosphere with 5% CO_2_ in air. Half-medium changes were performed at the end of each week with the same supplements, except for the omission of SCF and FLT3L after the first 2 weeks. At the end of week 3, cells from individual wells were harvested by trypsinization using trypsin+EDTA and collected into individual 1.2 mL tubes. After antibody staining, all samples were washed once and resuspended in HBSS containing 2% FBS, 200 μg/mL DNase and 1 μg/mL PI prior to their analysis on a flow cytometer. Detailed phenotypes for each lineage output are listed in Supplemental Table 2 and the limit of detection was set at 0.01% of total collected events with a minimum of 5 events detected.

### Xenotransplantation and analysis of regenerated human cells

Non-obese diabetic (NOD)-*Rag1*^−/−^*-Il2rg*^−/−^*-Kit*^W41/W41^ (NRG-W41) mice used in this study were bred and maintained in the Animal Resource Centre of the BC Cancer Research Institute. All animal experiments were performed according to the protocols approved by the Animal Care Committee of UBC. Sorted and unsorted CB populations containing 2-10 x 10^3^ CD34+CD38-cells were injected IV into each 7 to 10-week-old female NRG-W41 mouse immediately following exposure of the mice to 200 cGy of whole-body ^137^Cs γ radiation delivered over a few minutes, or 150 cGy of 300 KvP X-rays. At least 3 mice were transplanted in each experiment. Six to 13 weeks after transplantation, cells were collected from the BM of the engrafted mice. The RBCs in the samples were then lysed in cold Ammonium Chloride solution (STEMCELL Technologies) and the cells then passed through 40 μm filters and stained with antibodies in HBSS + 2% FBS, 1.5 µg/mL anti-human CD32 antibody and 10 µg/mL anti-mouse CD16/CD32 antibody. On the flow cytometer, human cells were identified using 2 anti-human CD45 antibodies with different epitopes. For subsequent functional analysis, the input cell subsets were sorted into Protein LoBind tubes (Eppendorf) or directly into tissue culture plates to initiate bulk and single-cell cultures, respectively.

### CFSE cell labeling and tracking

For CFSE staining, cells harvested from the LEM cultures or xenografts were washed twice in Dulbecco’s phosphate-buffered saline (PBS, STEMCELL Technologies) to remove residual serum. Cells were then resuspended in PBS containing 2.5 μM CFSE (STEMCELL Technologies) in a light-protected container and incubated at 37 °C in a water bath for 7 minutes. The cells were immediately washed 2 times, first with PBS + 10% FBS and then with αMEM + 7.5% FBS. Cells were then resuspended and incubated in the LEM culture condition overnight at 37 °C. On the next day, the cells were stained with antibodies and sorted on a flow cytometer. To reduce the heterogeneity of the input CFSE levels, the 30-40% of cells in the middle range of the CFSE peak were sorted^53^. For the “undivided” control, a small aliquot of sorted CFSE+ cells were treated with colcemid (Millipore-Sigma) to a final concentration of 0.25 μg/mL and then maintained under the same culture condition. Output cells were then collected at specified time points. To determine the division history of each phenotype being traced at a given time point, the levels of CFSE retention were measured by flow cytometry. The number of cell divisions completed was assigned based on a series of 2-fold decreases compared to the “undivided” control examined at the same time.

For CFSE tracking of cells produced in xenografts or from primary CB samples, cells were stained with CFSE using the steps described above, stored at 4 °C overnight to preserve cell viability and allow the diffusion of unbound CFSE molecules to reach an equilibrium, and then sorted the next day using the same procedures as above.

The average cell cycle transit time within each specified output phenotype at a given time point was calculated based on the numbers of divisions each cell has undergone since day 0. Proportions of cells that completed different numbers of divisions were converted to a series of probabilities. Z-scores were then derived from these probabilities assuming this follows a normal distribution. Mean and standard deviation values were then solved using the equation shown in Supplemental Fig. 3S using the two datapoints closest to the 50% probability within the distribution.

### Quantification of Ki67 and DAPI by immunofluorescence

P-mix cells were cultured in the LEM condition for 2 weeks and different output phenotypes were then isolated from the total by FACS. Each subset and freshly isolated CD34+38-CB cells were then transferred to separate wells in 384-well streptavidin-coated plates (Thermo Fisher) pre-treated with 1 μg/mL α-CD44-biotin (BioLegend)^54^. Cells were then fixed in 3% paraformaldehyde (Bio-rad) in PBS for 20 minutes at room temperature, and then washed twice with PBS + 2% FBS. Cells were then permeabilized with 0.5% Triton X-100 in PBS + 2% FBS for 20 minutes at room temperature, followed by a single wash with PBS + 2% FBS and then another with PBS + 5% FBS. Cells were then stained with 2.5 μg/mL anti-Ki67-AF488 (B56, BD Pharmingen) in PBS + 2% FBS for 16 hours at 4 °C. After 2 washes with PBS + 2% FBS, cells were resuspended in PBS+2% FBS supplemented with 1 μg/mL DAPI (Millipore-Sigma) for 15 min at room temperature and then imaged on a Nikon A1-Si Confocal microscope. The images acquired on the microscope were then analyzed using the ImageJ software. Selection of individual nuclei was performed using “threshold” and “create mask” functions. Integrated pixel intensities were then measured for each single nuclei. The results were further processed using the “flowCore” package in R. A size gate was first applied to remove any debris on the image. The intensities of Ki67 and DAPI in the nuclei were then plotted on a 2D plot where cells at different cell cycle phases were identified using the gates applied (see an example in Fig. 3g).

### Clonal liquid cultures and time-course phenotype tracking by fluorescence imaging

Twenty μL of LEM containing 160 ng/mL anti-CD34-AF647 (Biolegend), 50 ng/mL anti-CD45RA-BV421 (Biolegend) and 40 ng/mL anti-CLEC12A-PE (eBioscience) filtered through a 0.2 μm filter was dispensed into each well of a Terasaki plate (Greiner). Single input cells were sorted into each well of the Terasaki plates preloaded with the LEM culture medium. Each Terasaki plate was then placed inside a 15 cm culture dish together with 4-5 open 35 mm dishes containing sterile water to retain a sufficiently humidified atmosphere within the Terasaki plates. On Day 8, 3.5 μL of filtered fresh media with the same antibodies were added to each well of the Terasaki plates. The plates were imaged on a Nikon A1-Si Confocal microscope every 2 days from Day 2 to Day 16. Brightfield and fluorescent images of each well containing a clone were captured using a 10x objective that covered >95% of the bottom area of the Terasaki wells, with the x, y and z positions on the microscope adjusted for each well to include all of the output cells. Cell numbers of each clone were then counted on the brightfield images using the “Cell Counter” function in ImageJ. The fluorescent signals emitted from each surface antibody were used to define the phenotypes of the clonal outputs (listed in Supplemental Table 2).

A logistic growth model was used to fit the clonal expansion results using the R package “Grofit” and the equation shown below. Three parameters were derived from this model: the lag time (λ) defined as the inflection point where the starting flat line met the slope of the maximum growth phase; the maximum growth rate (μ) derived from the slope of the maximum growth phase; and the maximum growth (A) defined by the asymptote that the growth curve approached at the end. The “grofit” function was performed using the nonlinear least square fit algorithm where iterative model fitting was performed to reduce residual errors that converged on a final solution with a stable minimization of residual error. Individual clones were not fitted to the model if the algorithm failed to converge. The residual sum-of-squares, number of iterations and the achieved convergence tolerance were generated as a result to evaluate the qualities of the fitted models. The function “geom_curve” in the R package “ggplot2” was used to display the derived logistic functions on the growth curve plots.

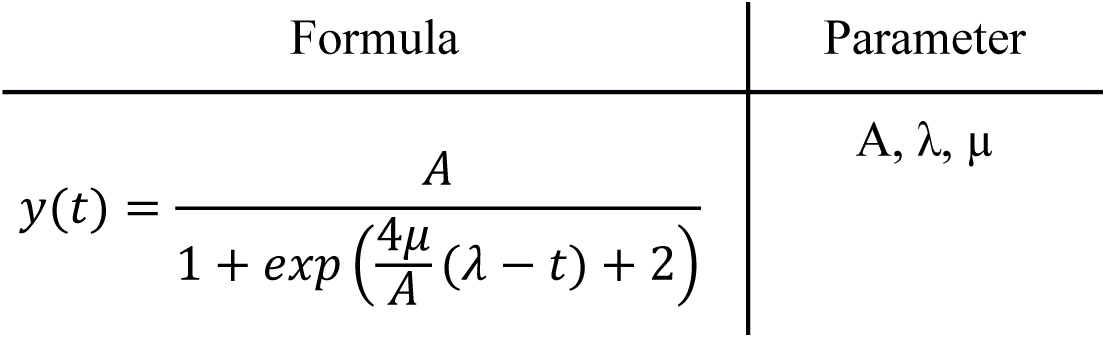

### RT-qPCR

P-mix cells were cultured in the LEM condition for 2 weeks and different output phenotypes were then isolated from the total output cells by FACS. To extract total RNA, 600-5,000 sorted cells were lysed in 250 µL TRIzol (Invitrogen). The lysate was then mixed first with 50 µL chloroform to release the RNA into the aqueous phase after centrifugation. 150 µL isopropanol was then mixed with the isolated aqueous phase and the samples incubated at −20 °C overnight. The precipitated RNA was then isolated by centrifugation, washed 2 times with 70% ethanol, and dissolved in RNase- and DNase-free water. Reverse transcription was performed using the Superscript VILO Master Mix (Invitrogen) according to the manufacturer’s instructions. The cDNA generated was then diluted 2-5 fold in RNase- and DNase-free water and mixed with PowerUP SYBR Green Master Mix (Applied Biosystems) and primers (listed in Supplemental Table 3) for input into a quantitative real time PCR reaction. Samples were then analyzed in a MicroAmp Optical 96-Well Reaction Plate (Applied Biosystems) using a 7500 Fast Real-time PCR Analyzer (Life Technologies). From this result, mRNA levels relative to GAPDH for each gene were calculated based on the differences of their C_t_ (threshold cycle) values.

### CITE-seq library generation and data processing

The input cells for the CITE-seq analysis were obtained from a pre-made pool of CB cells and processed following the 10x Genomics surface protein labelling protocol (CG000149). For the day 0 sample, a cryopreserved aliquot was thawed as described previously, but with omission of DNase I. Cells were then resuspended in PBS supplemented with 1% FBS and 9.1% TruStain FcX. Following a 10-minute incubation, cells were stained with both flow cytometry antibodies and TotalSeq antibodies (BioLegend) as follows: TotalSeq-B0853 anti-human CD371 (CLEC12A), TotalSeq-B0063 anti-human CD45RA, TotalSeq-B0062 anti-human CD10, TotalSeq-B0066 anti-human CD7, TotalSeq-B0054 anti-human CD34, TotalSeq-B0051 anti-human CD14, TotalSeq-B0392 anti-human CD15 (SSEA-1), CD45-AF700 (HI30, BioLegend), CD71-FITC (OKT9, eBioscience), CD38-PECY7 (HIT2, BioLegend), CD45RA-APC (MEM56, Invitrogen), CLEC12A-PE (HB3, eBioscience), CD10-BB700 (MEM78, BD Biosciences) and CD34-BV421 (561, BioLegend). Cells were then washed four times in PBS + 1% FBS before being sorted on a flow cytometer.

Samples for the day 7, 10, and 13 time points were obtained from cultured cells as shown in Fig. 6a. All samples were labeled with the same TotalSeq antibodies following the same protocol as for the day 0 cells, except for the day 10 and day 13 samples, which were not stained with flow cytometry antibodies. Additionally, the day 7 sample was stained with both flow cytometry and TotalSeq antibodies, albeit with a different panel for the former, including CD45-AF700 (HI30, BioLegend), CD45RA-APC (MEM56, Invitrogen), CLEC12A-PE (HB3, eBioscience), CD10-BB700 (MEM78, BD Biosciences), CD34-BV421 (561, BioLegend), CD14-PECY7 (61D3, eBioscience), and CD15-PECY7 (HI98, BD Pharmingen). After 1 hour, the day 7 cells were washed four times in PBS + 1% FBS before being sorted on a flow cytometer. Subsequently, the sorted (day 0, 7) and unsorted (day 10, 13) samples were resuspended to a final concentration of 800-1,000 cells/µL and processed at a sequencing facility in the SBME at UBC. Gene expression and surface protein libraries were generated using the Chromium Next GEM Single Cell 3 **’**Reagent Kits v3.1 and sequenced by Illumina MiSeq or NextSeq2000. Demultiplexed FASTQ files were processed using Cell Ranger version 7.1.0 (10x Genomics). Alignment of reads to the human reference genome GRCh38 and calculation of Unique Molecular Identifier (UMI) counts for gene expression and antibody libraries were performed using the “cellranger count” command. The UMI counts mapped to each cell were then compiled into a feature-barcode matrix. A total of 4,727, 3,419, 4,543 and 5,216 cells were initially detected by Cell Ranger for the day 0, 7, 10 and 13 samples, respectively. Across these 4 samples, the range of average reads per cell was 33,000-40,000 and the median number of genes detected was 3,700-5,300.

Gene expression analysis was done in R following the Bioconductor Single Cell Experiment pipeline (https://bioconductor.org/books/3.18/OSCA/). Quality control was performed using the “scuttle” package with a deviation-based threshold to filter out cells with low library size, low number of features and a high proportion of reads that mapped to the mitochondrial genome. Gene expression data were then normalized to library sizes and transformed to a log2 scale using the “scran” package. Doublet removal was performed using an in silico simulation method provided by the “scDblFinder” package. After the pre-processing steps, 4,088, 3,039, 4,252 and 4,890 cells remained for the day 0, 7, 10 and 13 samples, respectively. They were then pooled into a single dataset using the “multiBatchNorm” function to adjust for different library depths across samples. The top 30% variable genes were identified using the “modelGeneVar” function. The principal component (PC) analysis was performed on the top variable genes using the “runPCA” function. Further non-linear dimensionality reduction was then performed on the PCs using the “runUMAP” function with the number of components set at 2 or 4.

### Surface phenotype assignment

Quality control of antibody libraries was performed using the “DropletUtils” package. Cells with no detectable ambient contamination were removed. Normalization was then performed based on size factors derived from a simulated bimodal distribution of high expression signals and ambient contamination using the “ambientProfileBimodal” function in the same package. This was then followed by a log10 transformation of the normalized counts.

To correlate the measurements from ADTs to the phenotypes detected by flow cytometry, a proportion of each input sample used for CITE-seq was also analyzed on a flow cytometer. Flow cytometric data were then transformed using the asinh (x/150) function and different phenotypes were gated using the “flowCore” package. The scaled flow cytometric parameters were then used as the training data for the k-nearest neighbour classification. ADT data processed as above were also scaled and imported as the test dataset. The 10 nearest neighbours in Euclidean distance of the test data were then identified in the training data using the “knn” function of the “class” package. Cells were then assigned to the phenotype that was most frequently detected in their neighbours (ties broken at random).

### Cell cycle classification

The strategy for cell cycle segregation was based on the coordinated patterns of gene expression associated with G1/S and G2/M phases, respectively^55^. To create a first list of genes associated with different cell cycle phases, genes enlisted under the GO term “cell cycle” (GO:0007049) and previously identified in segregated phases of human myeloid leukemic cells^56^ were combined. This set was then intersected with the top 30% most variable genes from data pooled across all four time points. Additional cell cycle related gene expression patterns were searched in the principal component rotation vectors for each of the top 20 PCs and included in the final list (Supplemental Table 4). The “plotheatmap” function was used to generate a heatmap with hierarchical clustering across individual cells (columns) and individual genes (rows). Cells were divided into 6 clusters using the “cutree_cols” argument and then classified as G1/S, S or G2/M phases based on the expression of genes related to respective cell cycle phases, or as G0/G1 if they expressed all of these genes at low levels. Cell cycle regression was then performed using the “regressBatches” function in the “batchelor” package that removes a factor of variation associated with the 6 cell cycle clusters by fitting a linear model to the log-expression values for each gene and returns the residuals. Downstream analyses were then performed on the cell cycle-corrected residual values.

### Clustering of the CD34+RA-C-subset

Cells exhibiting the CD34+RA-C-phenotype within the total Cluster A population were isolated as a subset. Cell cycle regression was performed within the CD34+RA-C-subset using the method described above. Genes with a median expression of >0 before regression were selected for the PC analysis. The top 80 PCs generated were then used for a graph-based clustering method using the “clusterCells” function in the “scran” package. The parameters were configured with 25 nearest neighbors, the “Louvain” community detection method, and the “Jaccard” edge weighting scheme. Differential gene expression was analyzed using the “scoreMarkers” function from the same package. Highly expressed genes within each cluster were determined based on the mean area under the curve (AUC) value as an effect size, with a threshold set at >0.65. Enrichment of GO terms (biological process) was then performed on the identified marker genes using the “enrichGO” function from the “clusterProfiler” package with a q-value cutoff set at <0.05.

### Pseudotime trajectory modelling

For the trajectory analysis, cells within Cluster A were first grouped using the “clusterCells” function as described above. Lineage trajectories were then identified and fitted to principal curves embedded in a low-dimensional UMAP space using the “slingshot” package. Starting and ending clusters were not specified a priori. Pseudotime for each lineage path was computed using “slingPseudotime”, and genes exhibiting significant expression changes across pseudotime were identified by fitting a spline model, with its coefficients assessed through ANOVA tests via the “testPseudotime” function. To examine the expression dynamics of selected highly variable genes (FDR<1e-300) in more detail, cells along each lineage path were segmented into 10 equal-sized bins based on their pseudotime ordering, and the mean expression of each gene then calculated per bin. Genes were categorized as early, intermediate, or late-stage based on their peak average expression occurring within the 0-20%, 20-80%, or 80-100% intervals of the pseudotime distribution, respectively. Overlap among gene categories across different lineages was analyzed using the “UpSetR” package.

### Clustering of dynamically expressed genes in lineage trajectories

To identify gene expression patterns associated with different lineage outputs, genes exhibiting variable expression across distinct lineage trajectories were combined. Their variance was then analyzed using PCA, with individual cells considered as separate dimensions. The top 100 PCs were used for further dimensionality reduction through the “umap” function in the “uwot” package. The gene matrix and the dimensionality-reduced data was then imported into a SingleCellExperiment object. Genes were clustered using the “clusterCells” function, and each identified gene cluster was subjected to GO term enrichment analysis for biological processes, with a q-value cutoff <0.05.

### TF motif analysis using SCENIC

The SCENIC package was used to infer TF activities within the gene expression data, following the workflow developed for the R platform (https://htmlpreview.github.io/ https://github.com/aertslab/SCENIC/blob/master/inst/doc/SCENIC_Running.html). The input data comprised of log-transformed counts of the top 30% variable genes from cells collected on the day 7, 10 and 13 time points. Initial inference of positive or negative gene correlations was conducted using the Spearman correlation and “runCorrelation” function. Subsequently, co-expression between TFs and their putative target genes was determined using the “GRNBoost2” python package. The result from this co-expression analysis was then compiled into co-expression modules using the “runSCENIC_1_coexNetwork2modules” function with default parameters. Candidate TFs and target genes were then identified using the “runSCENIC_2_createRegulons” function. Motif information was derived from human databases that catalog motif rankings 500 bp upstream and 10 kb around the transcription start site (TSS) (data downloaded from “https://resources.aertslab.org/cistarget/databases/old/homo_sapiens/hg38/refseq_r80/mc9nr/gene_based/”). Motifs that obtain a Normalized Enrichment Score (NES) >3.0 are considered to be significantly enriched within the TF-target module, with only high-confidence motifs (either directly annotated or inferred through orthology) being preserved for subsequent analysis. A threshold for gene count was set at 15% of total genes for assessing the AUC of each gene set per motif ranking. The “iCisTarget” method was used to identify highly ranked genes for each motif. TF activities in individual cells were quantified using “runSCENIC_3_scoreCells”, producing binary outcomes (presence or absence) based on thresholds generated from the “AUCell_exploreThresholds” function. Binary heatmaps for selected TF motifs, most prevalent in specific lineages, were generated with the “plotheatmap” function from the “scater” package. TF motifs in day 0 cells were identified similarly using the steps above.

### Statistical analysis

Statistical analyses were performed using R software. All t-tests performed were 2-sided, and paired when appropriate using the function “t.test”. Comparisons between multiple groups were performed using the “pairwise.t.test” function with adjusted p-values calculated by the Holm correction.

## Acknowledgements

We thank M. Hale, G. Edin, A. Toner, Y. Y. M. Lau, and the staff of the Eaves Stem Cell Assay Laboratory for technical and project management assistance, including the acquisition and some of the initial processing of CB and BM samples. We thank T. Stach and her team at the sequencing facility at UBC SBME for their valuable advice on optimizing CITE-seq library generation procedures.

This work was supported by subawards to C.J.E. from the Terry Fox Foundation Program Project grant #1074 and the Canadian Cancer Society grant #705047. F.W. and L.G. held UBC Graduate Fellowships.

## Author contributions

F.W. and C.E. designed the experiments and wrote the manuscript; F.W. performed the *in vitro* and *in vivo* experiments and all data analysis. L.G. assisted with the xenograft experiments; C.H. and M.H. assisted with experimental design and interpretation; B.S. assisted with the statistical analysis of the B+NM clonal assays and interpretation; and all authors read and approved this manuscript.

## Data availability

In compliance with the institutional regulations regarding the publication of anonymized human samples, deposition of the raw data of the CITE-seq experiments to the European Genome-phenome Archive is being processed and anticipated to be accessible to registered users by Mar 31^th^, 2024.

## Code availability

Custom code will be made available on request.

## Supplemental figures

**Figure S1.**
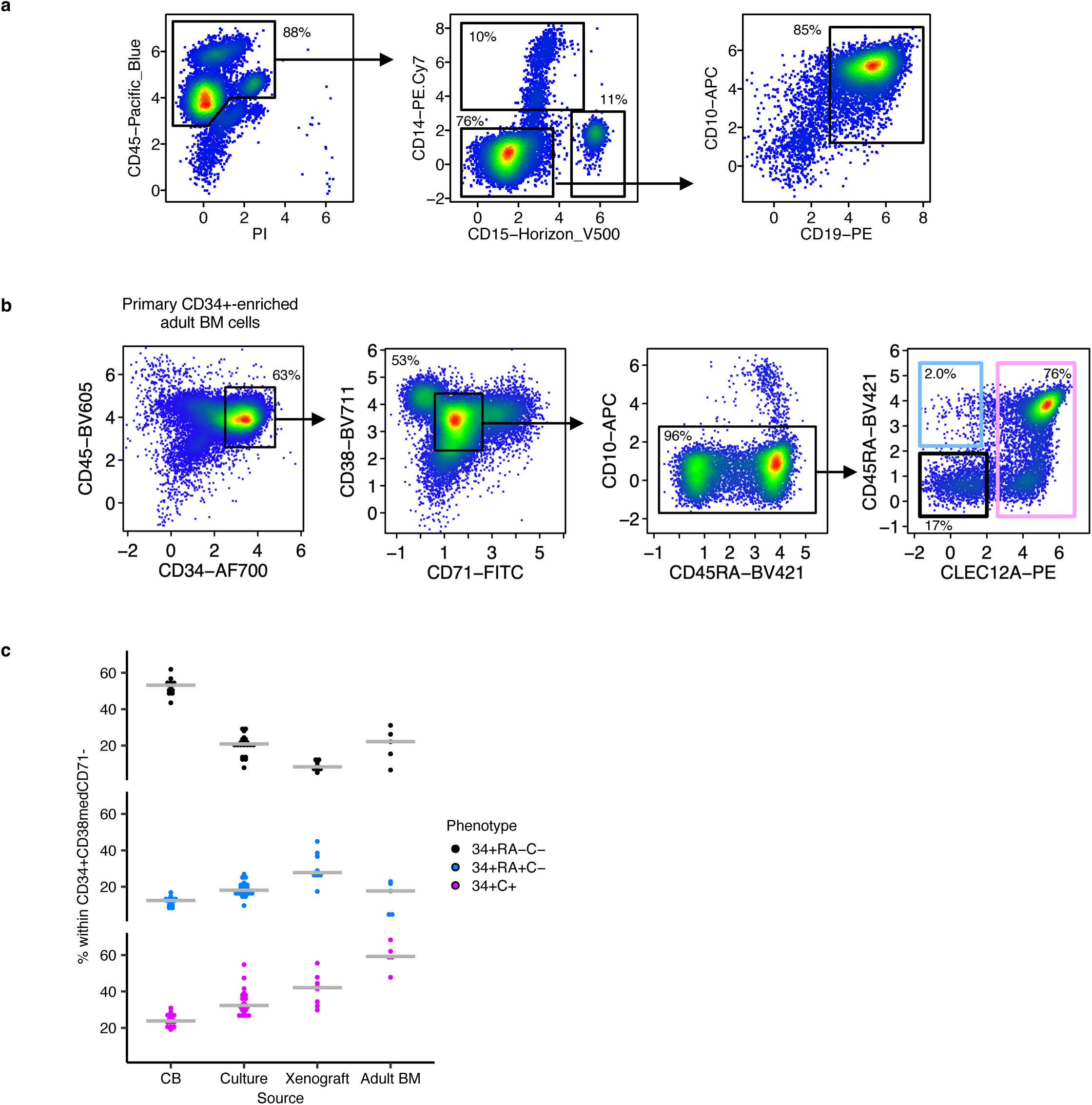
Phenotypic properties shared in hematopoietic cells from different sources. **(a)** An example of a bi-lineage clone produced in the clonal assay shown in Fig. 1d and the FACS gating used to define differentiated B and NM phenotypes. **(b)** An example of a flowcytometric profile of CD34+ cells isolated from a normal adult human BM sample. **(c)** Percentage of each of the CD34+RA-C-, CD34+RA+C- and CD34+C+ phenotypes in the CD34+38med71-fraction isolated from different tissue sources (each bar represents the median value).

**Figure S2.**
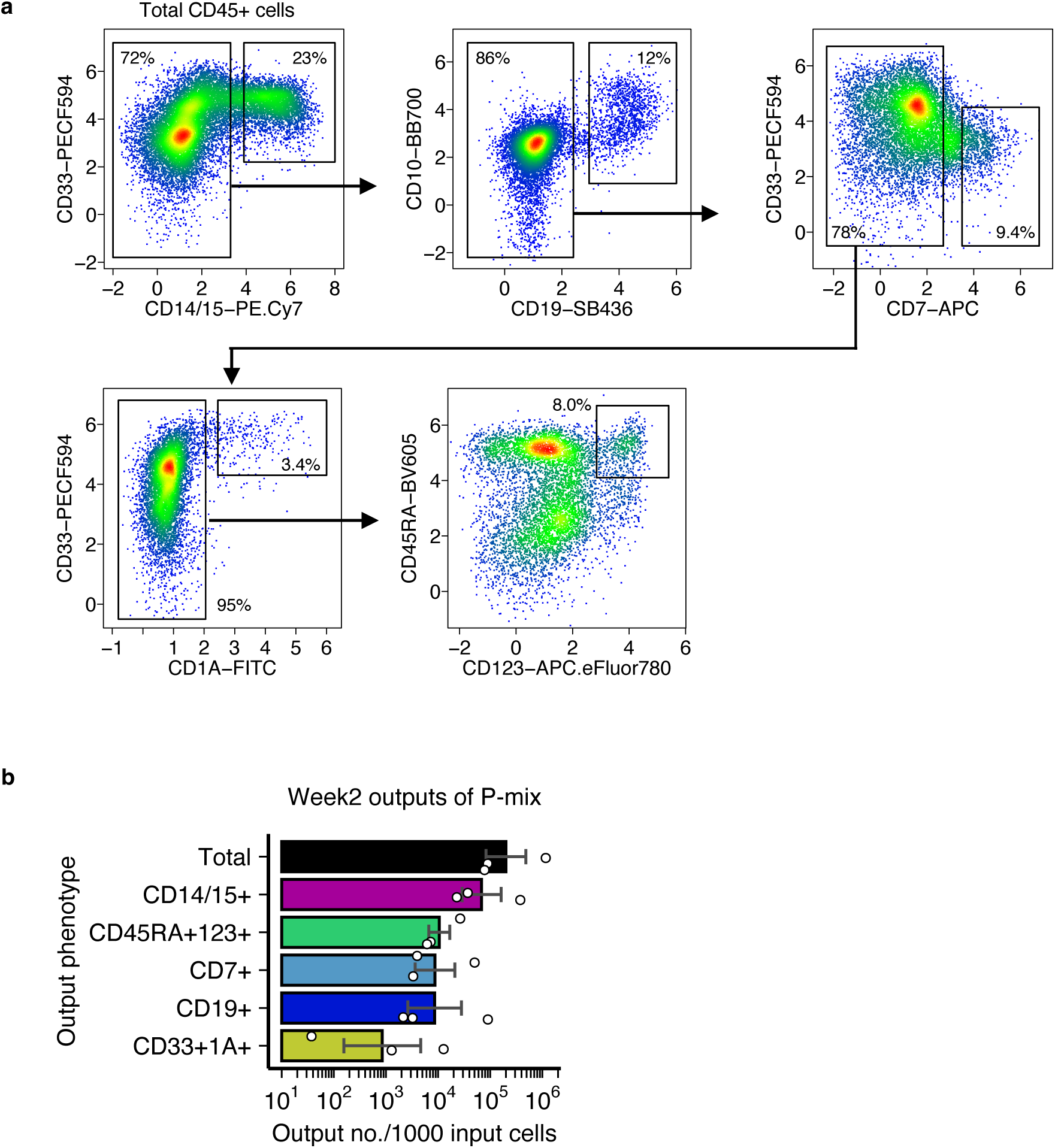
Gating scheme for other types of lineage precursors produced in the *in vitro* system. **(a)** An example of a flow cytometric profile of output phenotypes produced by P-mix cells analyzed at week 2 and the hierarchical gating for the NM, B, early-T, cDC and pDC lineage precursors, respectively. **(b)** Number of each output phenotype produced per 1,000 input P-mix cells analyzed at week 2. Each bar shows the mean ± SEM of values pooled from 3 different experiments.

**Figure S3.**
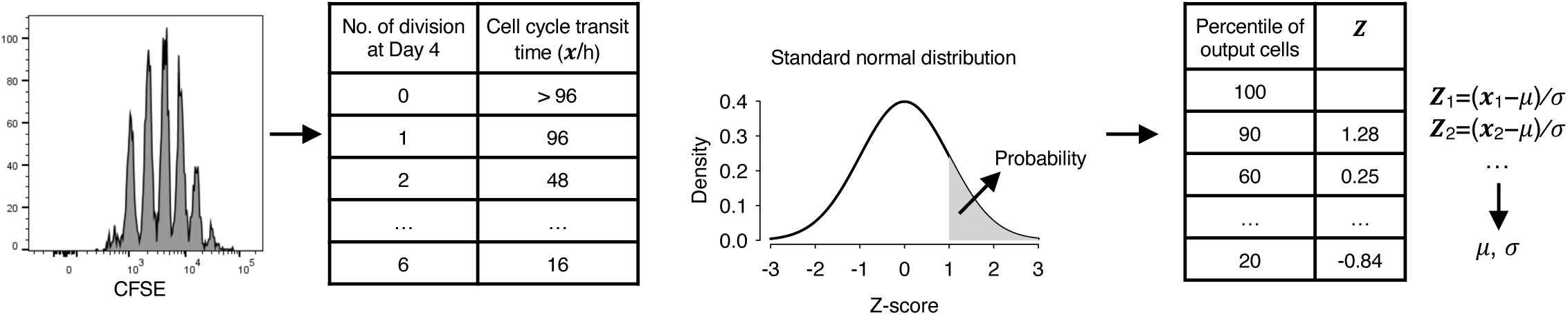
Description of the method used to calculate average cell cycle transit times from CFSE measurements using a day4 time point as an example.

**Figure S4.**
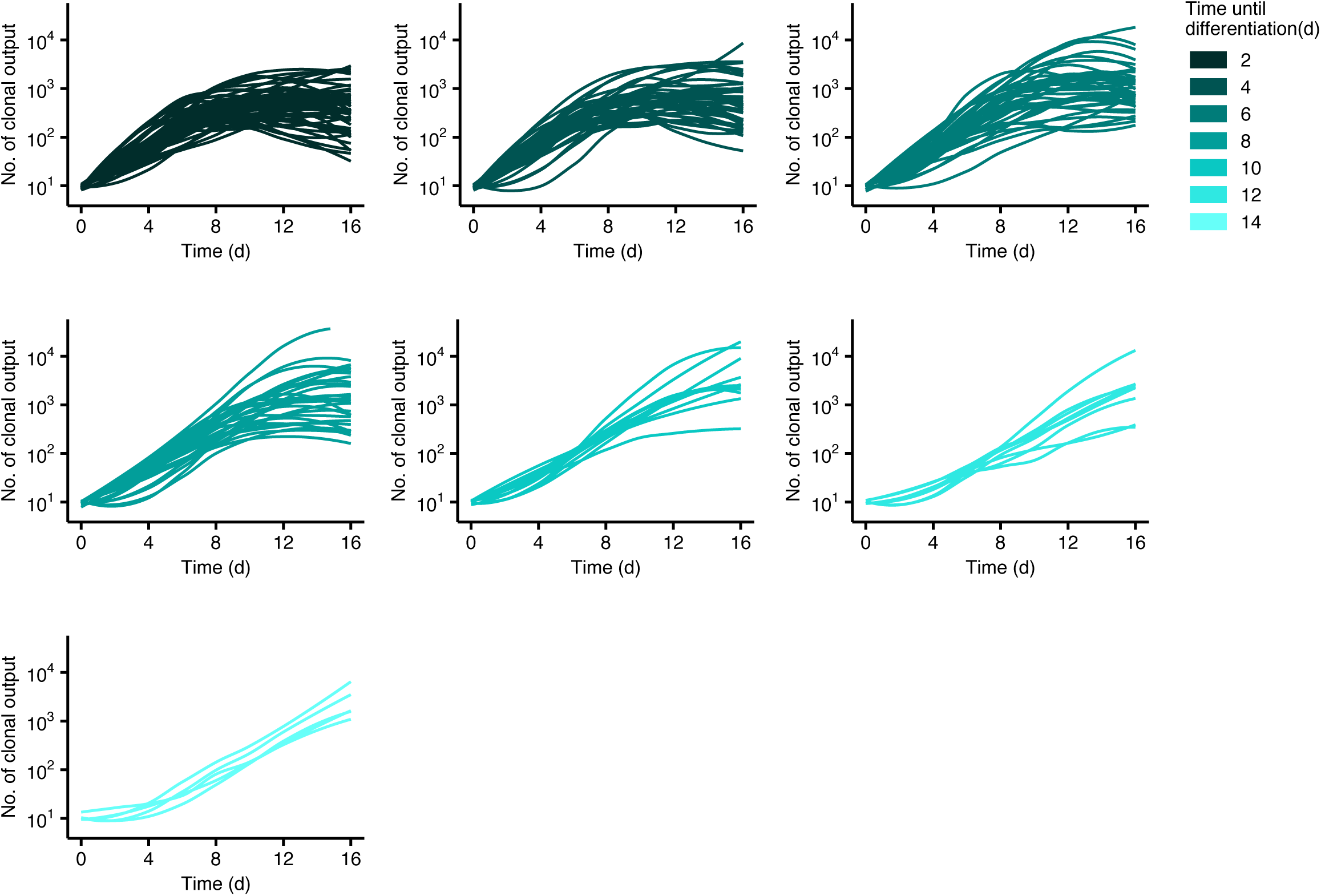
Overall growth kinetics (Lowess smoothed) of bi-lineage clones in each subgroup defined in Fig. 4e.

**Figure S5.**
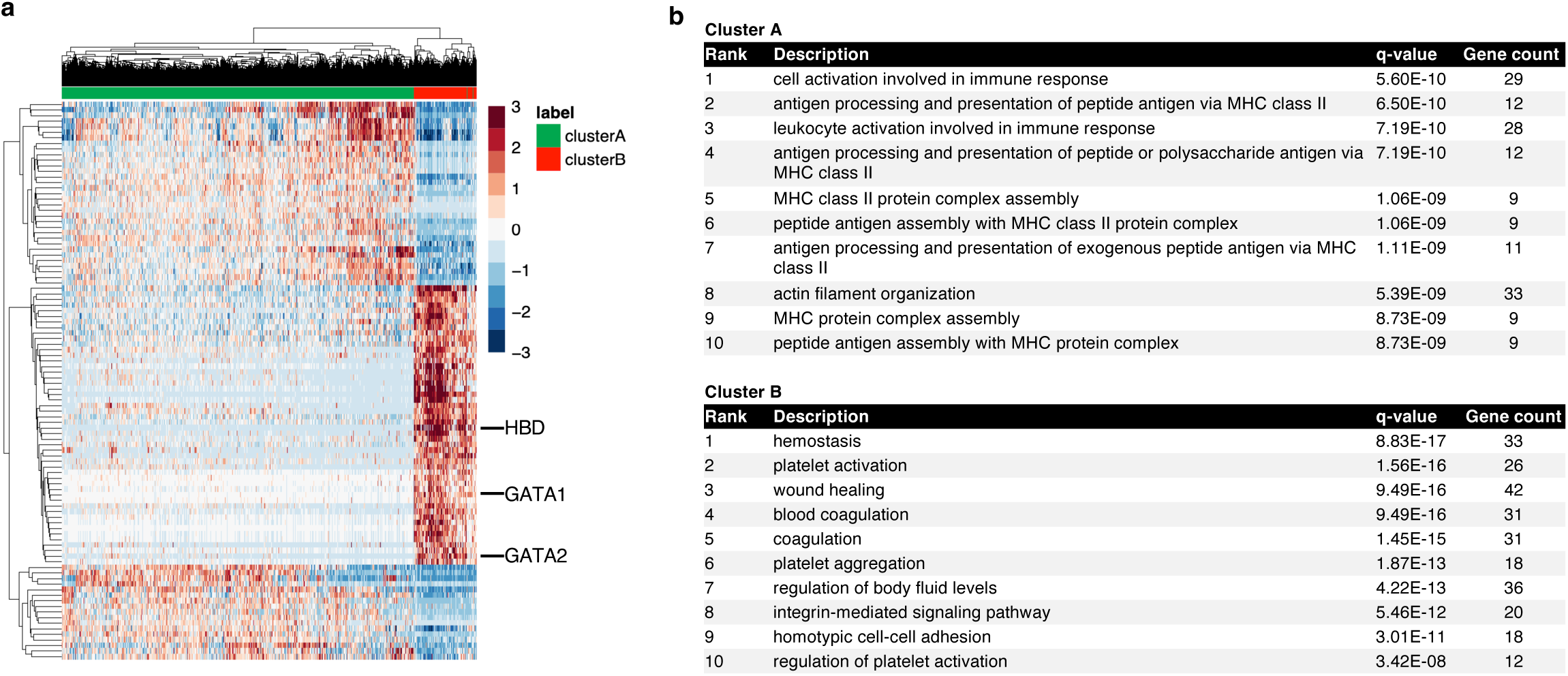
Marker genes highly expressed in Cluster A and Cluster B cells. **(a)** Hierarchical clustering of the 12,181 single cells pooled from day 7, 10 and 13 time points based on the top ranked differentially expressed genes between Cluster A and B cells defined in Fig. 6c. **(b)** Top 10 GO terms enriched in differentially expressed genes identified from Cluster A and B respectively (q-value cutoff <0.05).

**Figure S6.**
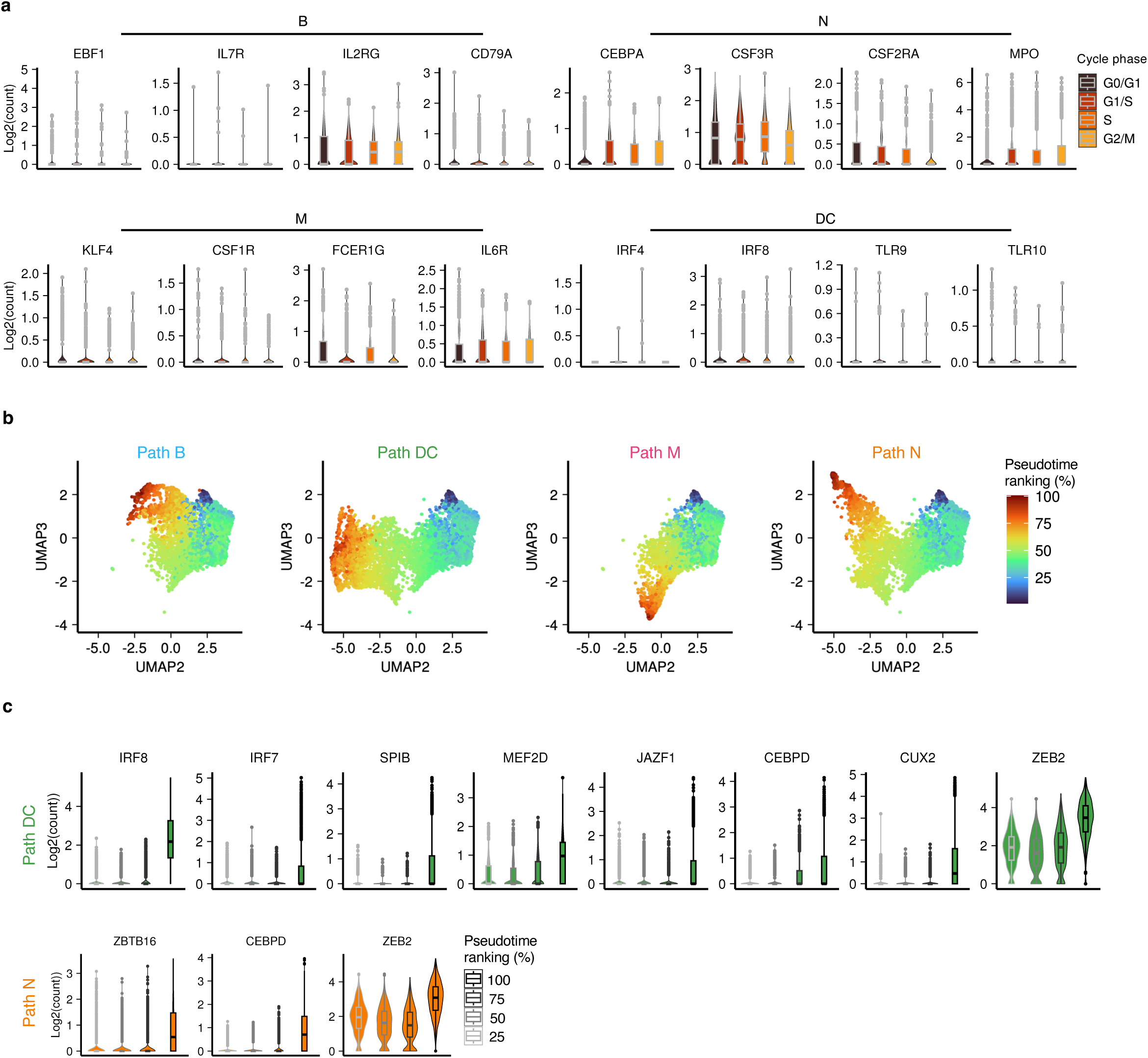
Dynamic gene expression changes during lineage restriction processes. **(a)** Log-transformed transcript counts of genes historically associated with specific lineage functions detected in cells assigned to different cell cycle phases (center line=median; box limits=first and third quartiles; whiskers=1.5x interquartile range; points=outliers). **(b)** UMAP presentation of cells belonging to each trajectory path. Cells have been coloured according to their pseudotime ranking. **(c)** Expression of “late” genes in the DC and N trajectories. Cells were grouped by the quadrants of their pseudotime ranking (center line=median; box limits=first and third quartiles; whiskers=1.5x interquartile range; points=outliers).

**Figure S7.**
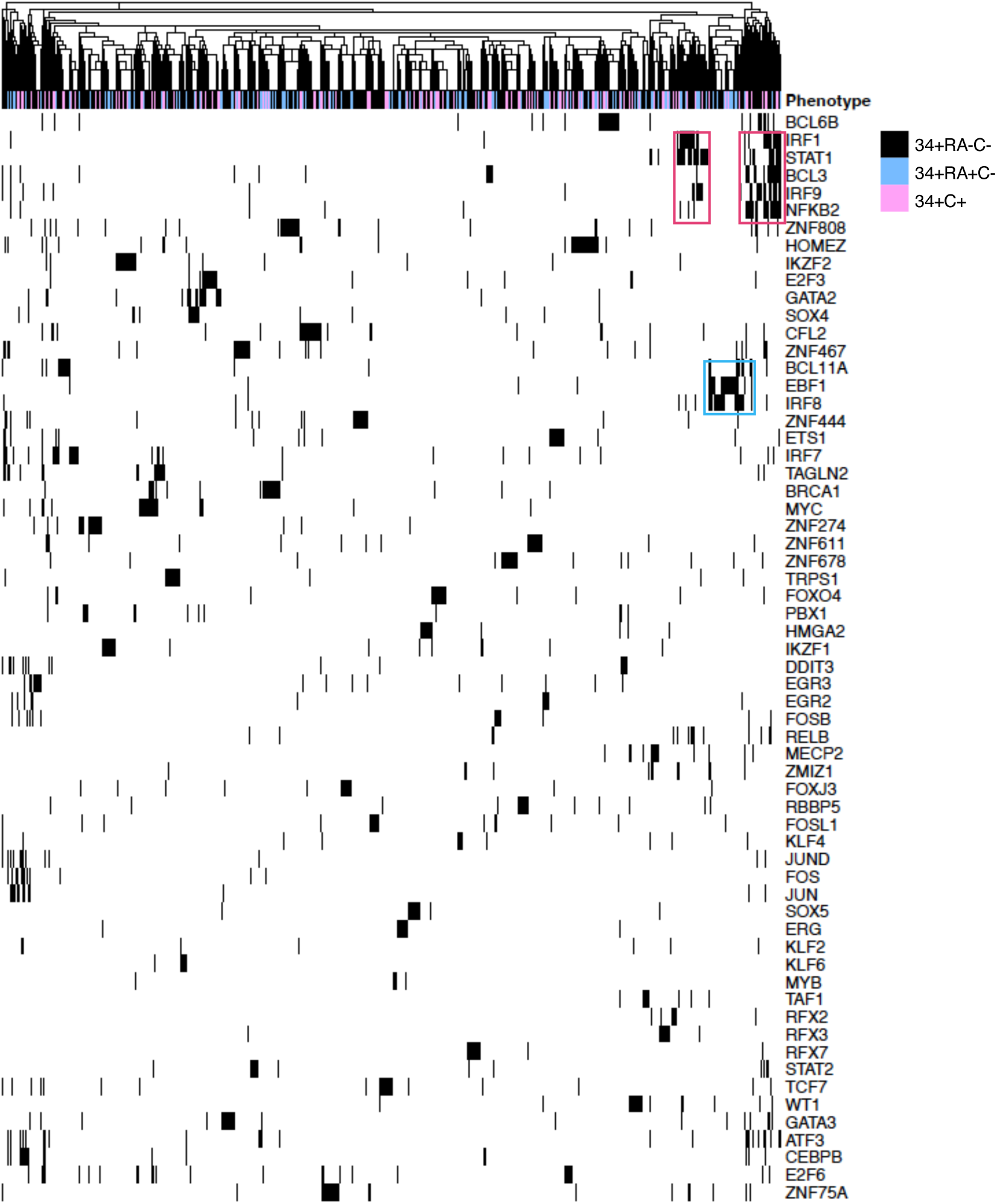
TF motif distribution in primary P-mix cells. Status of 68 TF motifs identified in day 0 P-mix cells, each bar indicating the presence (black) or absence (white) of each motif in individual cells. Columns are coloured by the phenotype of each cell.

